# *RAI1* safeguards fidelity and tempo of human neurodevelopmental gene expression

**DOI:** 10.64898/2026.06.11.731717

**Authors:** Bo Zhou, Satabdi Mohanty, Paris Riggle, Takao Tsukahara, Grace Lin, Louis T. Dang, Michael A. Sutton, Shigeki Iwase

## Abstract

Human brain development proceeds on an unusually long timeline relative to other species, a feature that is thought to foster advanced cognitive abilities. The Retinoic Acid Induced 1 (RAI1) gene encodes a nucleosome-binding protein, and its haploinsufficiency is responsible for Smith–Magenis Syndrome (SMS), a neurodevelopmental disorder characterized by cognitive impairment with autistic features. However, the role of *RAI1* in human neurodevelopment remains unexplored experimentally. Here, we generated isogenic heterozygous and homozygous *RAI1* loss-of-function human embryonic stem cell lines and interrogated the roles of *RAI1* in neurodevelopmental gene regulation. A longitudinal transcriptome analysis during *in vitro* cortical development revealed that *RAI1* deficiency accelerates the progression of developmental gene expression. Single-cell RNA-seq analysis revealed that *RAI1-*deficient neuroprogenitors acquire a transient mesoderm-like gene expression signature, followed by a pro-neuronal maturation signature in postmitotic neurons. Unexpectedly, the developmental acceleration signature was exacerbated during NGN2-induced excitatory neuron differentiation, isolating the roles of RAI1 in neuronal differentiation from non-neuronal functions. Together, these results identify *RAI1* as a suppressor of the mesodermal lineage program and as a brake that slows the tempo of human neurodevelopmental gene expression.

## Introduction

Human brain development unfolds over a longer timeline than in other mammals even when normalized to their lifespans. While rodent neurons achieve functional maturity relatively soon after birth, human cortical neurons require years to complete their developmental program, with an extended period of neural progenitor cell (NPC) expansion followed by synaptic refinement continuing into adolescence and beyond (1,2). This extended timeline leads to developmental neoteny and is thought to enable the expanded learning capacity and behavioral flexibility that characterize human cognition (1,3). While possibly advantageous for higher cognitive capacity, developmental neoteny may make the human brain vulnerable to neurodevelopmental abnormalities such as autism (3,4). Thus, determining how species-specific developmental timing arises is essential for understanding not only normal brain development but also the pathogenesis of neurodevelopmental disorders.

Genetic mechanisms underlying the extended timeline of human neurodevelopment have begun to emerge in the past decade. Changes in expression timing of conserved genes such as *ZEB2* help drive human-specific traits such as the pronounced expansion of radial glia cells (5). Human- and primate-specific genes that emerged through duplication and diversification also contribute. *NOTCH2NL* genes, which arose from a partial duplication of *NOTCH2*, support the expansion of radial glia cells, and *SRGAP2C*, a counteracting homolog of the evolutionarily conserved *SRGAP2A*, promotes dendritic spine maturation (6). In addition to genetic mechanisms, metabolic changes such as the slow transition from glycolysis to oxidative phosphorylation during neuronal differentiation contribute to the extended timeline (7). Thus, diverse mechanisms are at play to slow neurodevelopment, fostering higher cognition in humans.

More recently, a group of evolutionarily conserved chromatin regulators has been shown to extend the maturation period of human neurons. These chromatin regulators include histone methyltransferases such as EZH2 (H3K27), EHMT1 and EHMT2 (H3K9), and DOT1L (H3K79), histone demethylases such as KDM1A and KDM5B, and ATP-dependent chromatin remodelers such as SMARCE1 and CHD3 (8). Expression of these chromatin regulators is high in immature excitatory neurons but is downregulated during the synaptogenesis phase of in vitro differentiation of pluripotent stem cells (hiPSCs). When genes were deleted at NPC stages, neuronal maturation was significantly accelerated. In mouse neuronal differentiation, *Ezh2* expression was lower than in humans, and murine *Ezh2* depletion led to a mild upregulation of neurodevelopmental gene expression. Thus, the roles of *Ezh2* in suppressing neurodevelopmental gene expression programs appear conserved across species, with its higher expression in human neurons conferring protracted neurodevelopment. However, it remains unknown whether the current list of chromatin genes that slow human neurodevelopment is complete.

Chromatin regulators, when mutated, are a major genetic driver of neurodevelopmental disorders (1,9–11). Among such genes is Retinoic Acid Induced 1 (*RAI1*), which encodes a nucleosome-binding protein associated with Smith-Magenis Syndrome (SMS), a neurodevelopmental disorder characterized by intellectual disability, self-injurious behavior, obesity, and sleep disturbances (12). While the majority of SMS cases arise from heterozygous deletion of the 17p11.2 interval, spanning up to 50 genes, ∼ 10% of patients have heterozygous loss-of-function of *RAI1* alone (13). These patients exhibit SMS features similar to those with 17p11.2 deletions, thus leading to a consensus that *RAI1* is the key disease gene in the interval, though other genes may impact full SMS pathophysiology (14). SMS mouse models of *RAI1* deficiency recapitulate many SMS features, including obesity and learning deficits (15–18). However, certain phenotypes, such as ventriculomegaly—enlarged ventricles —seen in some SMS individuals are absent in SMS mouse models (19).

A recent report using induced pluripotent stem cells (iPSCs) from SMS individuals with 17p11.2 deletions reported accelerated neuronal differentiation, premature cell-cycle exit of neural progenitor cells (NPC), and increased intrinsic excitability (19). The premature cell-cycle exit observed in iPSC-derived NPCs may explain the ventriculomegaly observed only in human SMS patients. Since most studies analyzing mouse models of SMS used targeted *RAI1* deletion, and the human study involves a 17p11.2 deletion spanning dozens of genes, it remains unknown whether the phenotypic differences between species are due to species-specific roles of *RAI1* or other genes in the deleted interval. As such, the role of *RAI1* specifically in human neurodevelopment remains unclear.

In the present work, we generated *RAI1*-deficient human embryonic stem cells (hESCs) and examined the roles of *RAI1* in human neurodevelopment and gene expression. The use of hESCs enables us to examine the impact of *RAI1* loss relative to isogenic controls, unlike prior work with patient-derived iPSCs. We find that targeted loss of *RAI1* in human neurons is sufficient to accelerate neurodevelopmental gene expression programs and lead to faster maturation of excitatory synaptic function. No such features are evident in mouse transcriptomic data, suggesting this role may be species-specific. Our results establish *RAI1* as a new chromatin regulator that contributes to protracted human neurodevelopment. In addition, our data indicated a previously undocumented role of RAI1 in suppressing the mesodermal gene expression program during early development.

## Results

### Generation of *RAI1* mutant human embryonic stem cells

To investigate the role of *RAI1* in human neurodevelopment, we generated heterozygous (+/-) and homozygous (-/-) *RAI1* knockout lines in H9 human embryonic stem cells (hESCs) using CRISPR/Cas9 gene editing (Fig. 1A). A guide RNA was designed within Exon 3, and CRISPR-induced single-nucleotide insertion and deletion were observed in heterozygous and homozygous clones, respectively. We chose 2 independent clones with identical indels for *RAI1*+/-(HET15, HET18) and *RAI1*-/-(SC6, SC9) for further study. SNP microarray analysis did not detect chromosomal abnormalities during gene editing (Fig. 1B). The introduced mutations caused frameshifts in both heterozygous and homozygous mutants. The premature stop codons did not result in nonsense-mediated decay (NMD), likely due to the length of the targeted exon 3 (20) — 1,701 bp encoding the majority of RAI1 protein—judged by the comparable read counts in bulk mRNA-seq (Fig. 1C). Western blots, using an in-house RAI1 antibody recognizing the N-terminus (21), validated the expected loss/reduction of RAI1 protein in mutant cells, without detecting a frame-shift-induced truncated protein (Fig. 1D). Similar truncating mutations have been found in SMS patients (22). Immunofluorescence analysis of pluripotency markers, including NANOG, OCT3/4, SOX2, and SSEA4, demonstrated their normal expression and ESC morphology (Fig. 1E).

**Figure 1.**
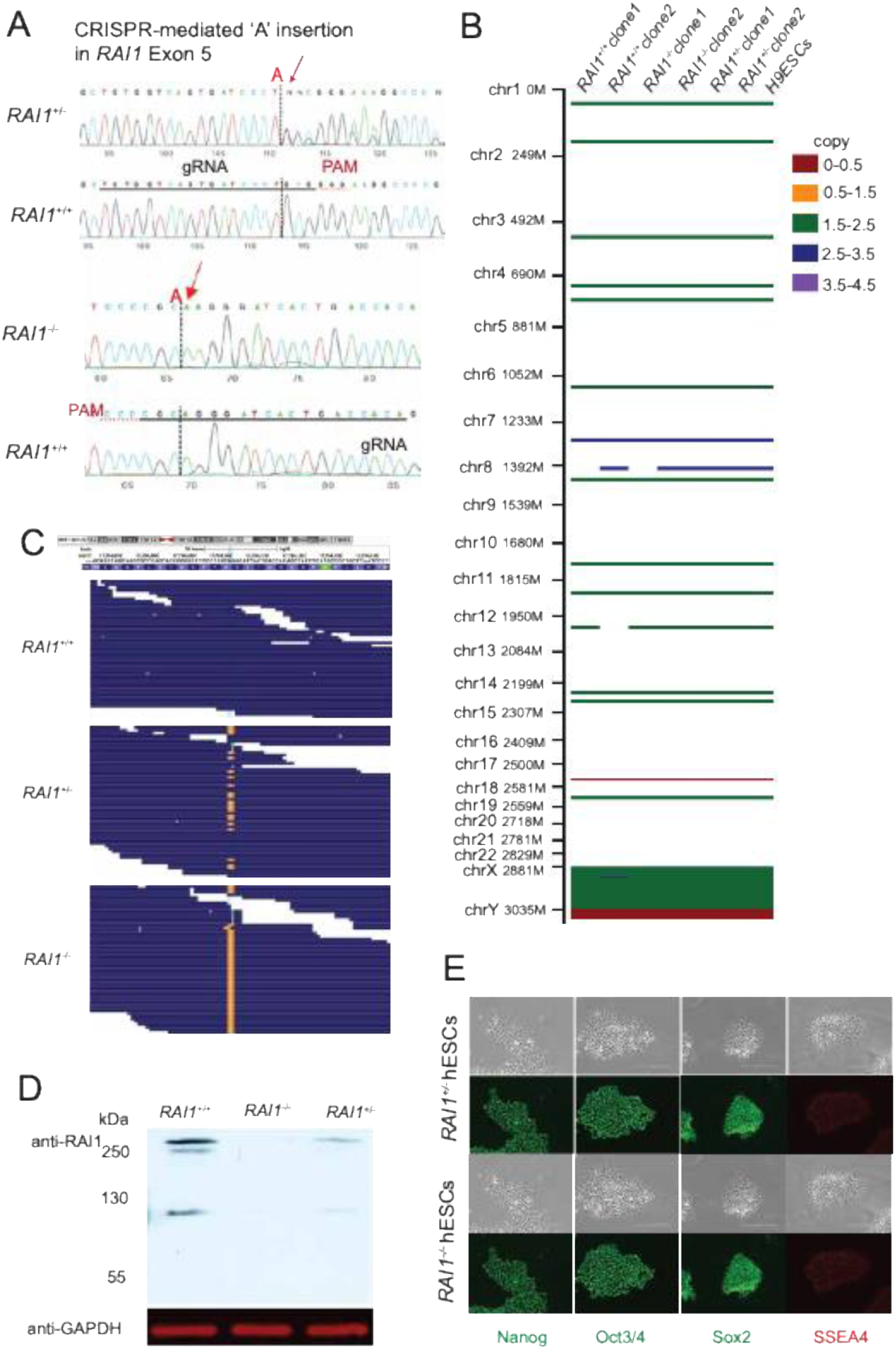
Generation of *RAI1*-deficient hESCs. **(A**) CRISPR gRNA design resulted in indels, and Sanger sequence validation. Note: Sanger sequence shows the opposite strands of the same target sequence for heterozygous and homozygous hESC lines. **(B)** Copy number measurements with SNP microarray. Copy numbers are summarized. White color denotes no copy number change compared to hg19. **(C)** Bulk mRNAseq reads demonstrating alignment mismatches (orange) in heterozygous and homozygous mutant cells (DIV20) due to the CRISPR-generated indels. **(D)** Western blot analysis of *RAI1* in iNeurons (DIV 7th) derived from mutant and WT hESCs. GAPDH: loading control. **(E)** Morphology of undifferentiated hESCs and immunofluorescence of pluripotency markers.

### Neurodevelopmental gene expression of *RAI1*-deficient cells

We next differentiated the hESC lines into excitatory neurons with a dual SMAD inhibition protocol (23) (Fig. 2A). To track the developmental trajectory of gene expression changes, we collected RNA from day 0, 20, 34, and 60 of differentiation and performed bulk mRNA-seq. *RAI1* expression increased continuously throughout differentiation. Principal component analysis (PCA) delineated the progression of transcriptomic changes over the 60 days. The transcriptomes of *RAI1*-homozygous mutants exhibited a noticeable acceleration of differentiation signatures, most pronounced at day 34, whereas RAI1-heterozygous mutant cells did not show a consistent difference across time points (Fig. 2C and D). The *RAI1*-heterozygous and homozygous mutants had comparable numbers of differentially expressed genes (DEGs, Padj < 0.01, Log2FC > 2, list: Table S1). DEGs in heterozygous and homozygous mutants overlap significantly, while some DEGs appear only in one mutant (Fig. S2). The number of DEGs was much greater in differentiated neurons than in undifferentiated hESCs (day 0), indicating a greater role for RAI1 in differentiation than in stem cells.

**Figure 2.**
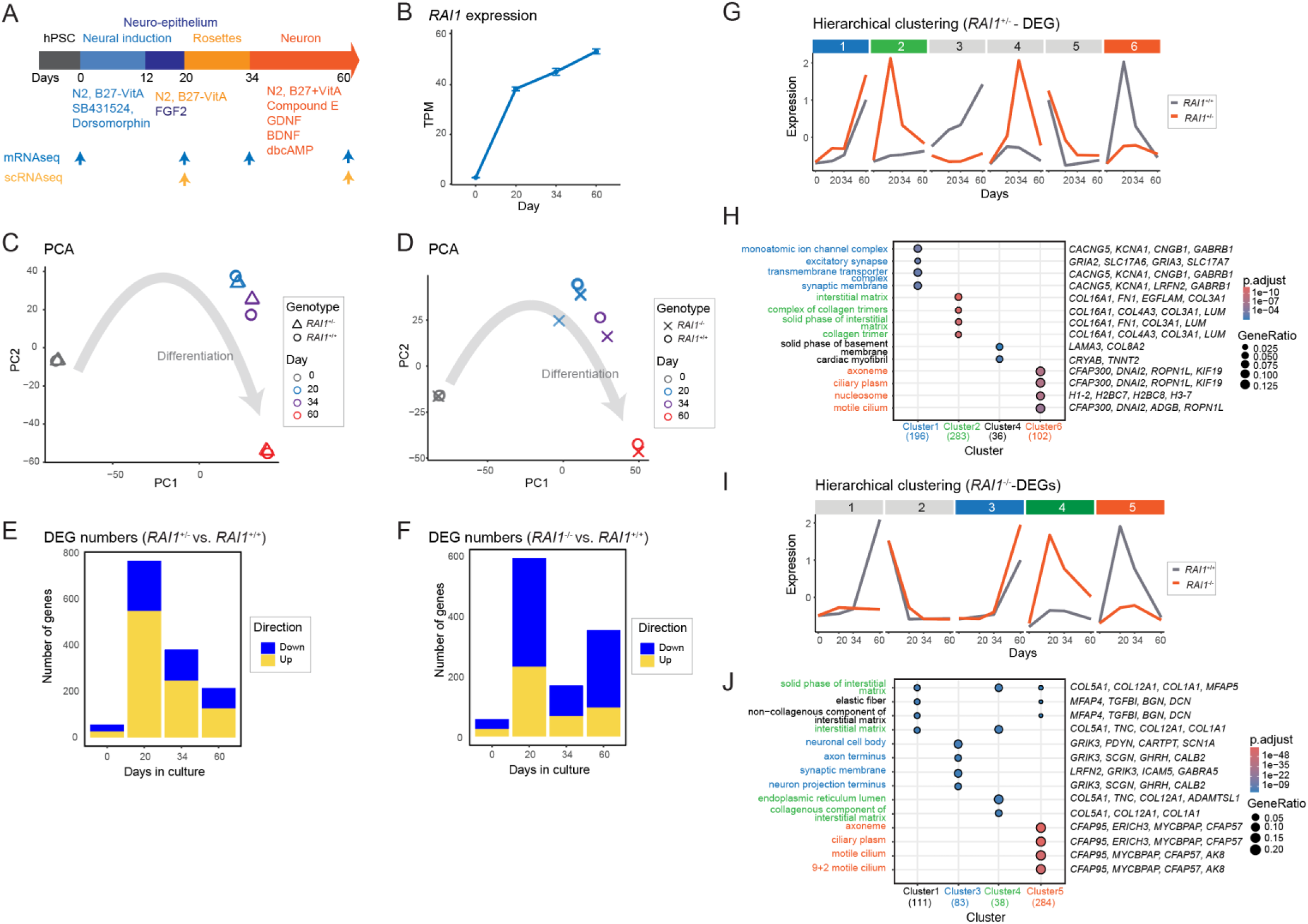
The time-series mRNA-seq revealed acceleration of the neurodevelopmental gene expression program in *RAI1*-KO neurons. **(A)** Schematics for neuronal differentiation and RNA-seq experiments. **(B)** RA1 expression from the mRNA-seq data. **(C&D)** PCA analysis of WT vs mutant (C: heterozygous, D: homozygous) mRNA-seq data. (n=4, per genotype). **(E&F)** The number of DEGs (E: heterozygous, F: homozygous, |Log2FC| > 2, Padj < 0.01). **(G&I)** Hierarchical clustering of all DEGs (G: heterozygous, I: homozygous). Line plots: median Z-scores of each cluster. **(H&J)** Gene ontology enrichment of DEG clusters (Cellular components, H: heterozygous, J: homozygous). The top 4 genes contributing to ontology enrichment are indicated.

To gain insights into biological processes impacted in the mutants, we performed unbiased hierarchical clustering of all DEGs followed by gene ontology enrichment analysis (Fig. 2. G-J). The clustering captured three gene groups that show consistent changes in heterozygous and homozygous mutants during differentiation, and the three clusters were enriched for similar or identical GO terms (colored in Fig. 2. G-J). These are (1) synaptic genes: accelerated expression in the mutants (blue), (2) collagen-related genes: ectopic induction at days 20 and 34 (green), and (3) cilia-related genes: failed induction at days 20 and 34 (orange). Taken together, *RAI1* loss, both heterozygous and homozygous, leads to noticeable acceleration of synaptic gene expression, accompanied by dysregulation of collagen- and cilia-related gene expression during dual SMADi-induced neuronal differentiation.

### The impact of *RAI1* loss on cell types and cell cycle phases

The above bulk RNA-seq results showed clear changes in gene expression; however, these changes may have reflected either cell-type composition or mRNA abundance within a given cell type. To distinguish between these possibilities, we performed single-cell RNA-seq (scRNA-seq) at day 20 and day 60 of dual SMADi differentiation (Fig. 1A). Data were processed with Seurat (24); a total of 100,191 cells passed the quality controls, including low mitochondrial read counts, and the expression of 308-12,372 genes was detected in individual cells with more than 1 read. Two replicates from independent hESC clones showed highly concordant UMAP profiles (Fig. 3A). The chosen resolution gave 7 clusters identified as neural progenitor G2/M (NPC-G2M), NPC-G1, intermediate progenitor cells (IPC), neuron, mesenchymal wild-type (Mes-WT), mesenchymal KO-type (Mes-KO), and gliogenic radial glia (gRG), based on marker gene expression (Fig. 3B) and inferred cell cycle phases (Fig. 3E). *RAI1* was expressed in all cell types at similar levels, with the highest expression in neurons (Fig. 3C).

**Figure 3:**
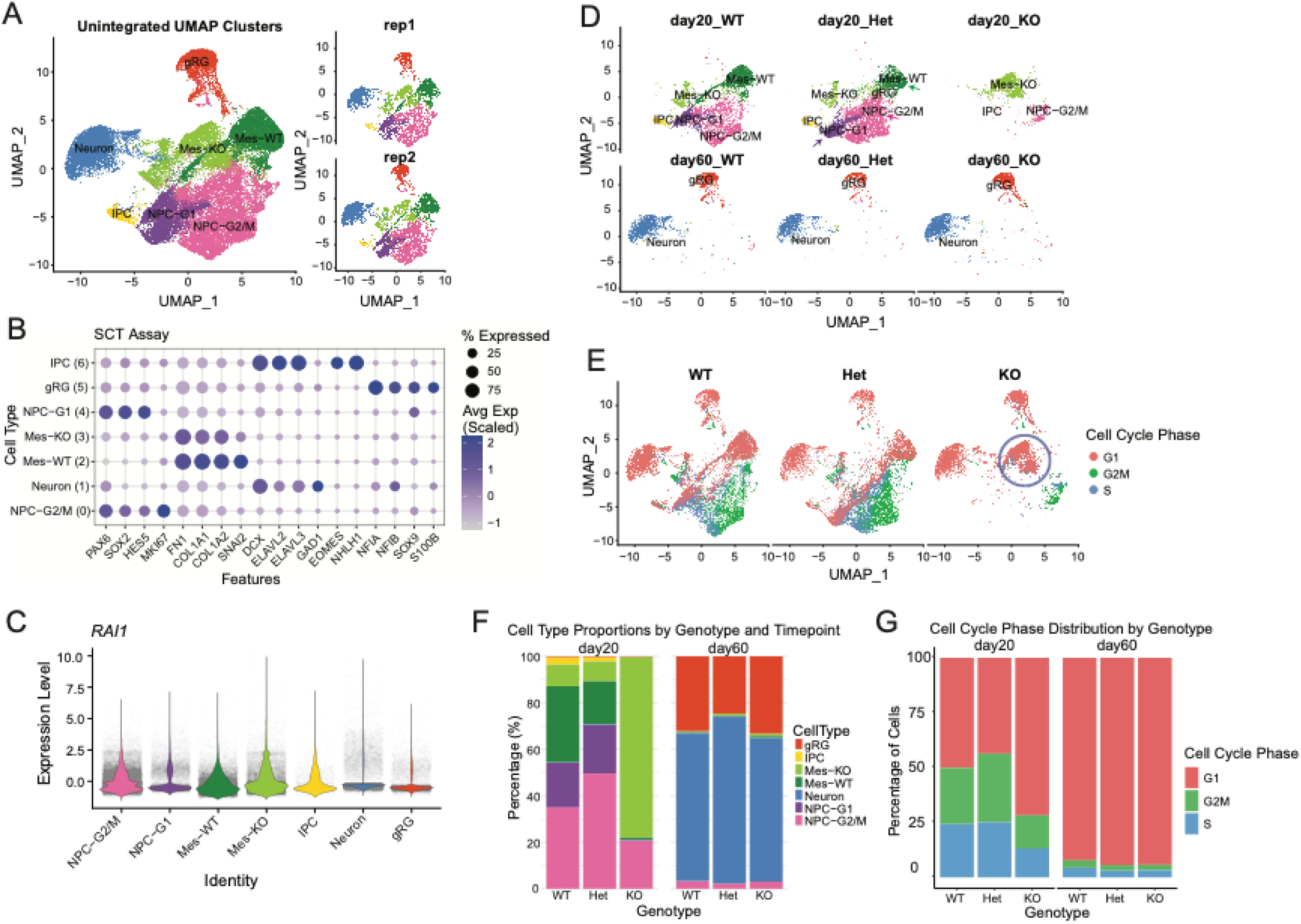
scRNA-seq: cell type composition and cell cycles in *RAI1*-deficient cells. **(A)** Uniform Manifold Approximation and Projection (UMAP) plot of all data combined identified 7 major clusters. **(B)** Marker gene expression used for the cell type inference of the 7 clusters. **(C)** *RAI1* expression in each cell type. **(D)** UMAP plots separated by genotypes and time points. **(E)** Cell cycle phases determined by gene expression. **(F)** Cell numbers of each cell types across genotypes. **(G)** Cell numbers of each cell cycle phase.

At day 20, the WT culture consisted of NPCs, IPCs (early neurons), and mesenchymal cells, and at day 60, differentiated neurons and gliogenic radial glia (gRG) appeared (Fig. 3D). This pattern was markedly disrupted in *RAI1*-homozygous mutants at day 20, where NPC-G1 and IPC were nearly absent, and the transcriptomic profiles of mesenchymal cells clearly shifted, hence its name: Mes-KO (Fig. 3D & F). The UMAP profile of the *RAI1* heterozygous mutant at day 20 was overall similar to WT; however, it showed noticeable differences in the NPC-G1, NPC-G2/M, and Mes-WT populations, suggesting changes in gene expression and cellular states (Fig. 3D, colored arrows). At day 20, *RAI1* heterozygous cultures showed a 21% increase in NPC at the expense of other differentiated cell types (Fig. 3F). Despite these large changes in cell populations at day 20, day 60 UMAP profiles were similar across genotypes except for a slight increase in neurons in the heterozygote. Cell cycle marker expression showed that approximately half of the WT cells were dividing at day 20, with most exiting the cell cycle by day 60 (Fig. 3E&G). Meanwhile, ∼75% of *RAI1* homozygous mutant cells had already entered G1 phase at day 20 (blue circle in Fig. 3E and Fig. 3G). The impact of *RAI1*-heterozygosity was milder, yet they showed a 7% increase in NPC-G2/M at the expense of G1 cells (Fig. 3F), suggesting a slightly delayed differentiation. These cell type composition and cell cycle phase analyses collectively suggested that (1) some RAI1 homozygous cells precociously differentiated into mesenchymal cells, (2) RAI1 heterozygous mutants underwent a delayed differentiation of NPCs at day 20, (3) however, eventually at day 60, all genotypes could produce neurons and gRG.

Mesenchymal cells in this differentiation protocol are likely neural crest cells, because their emergence relies on WNT signaling (25), and our differentiation protocol did not include a WNT inhibitor. This population was abundant on day 20 but nearly absent on day 60, suggesting that neural crest cells appeared early in the differentiation process and were eliminated over time due to culture conditions optimized for excitatory neuron survival. The compositional and transcriptomic changes in this cell population highlight a previously uncharacterized role for *RAI1* in neural crest cells.

### *RAI1* loss leads to accelerated differentiation of both neurons and mesenchymal cells

Next, we sought to determine if the altered developmental tempo of *RAI1*-deficient cells is cell-type specific. We used Monocle 3, an unsupervised algorithm, to infer the temporal order of single-cell transcriptomes (26). The Monocle 3 trajectory map largely corresponded to Seurat cell types (Fig. 4A). We set the dimensionality to 5 to prevent clusters from completely separating and calculated pseudotime for all cells using single roots: 10 randomly selected cells from the WT NPC-G2/M population. The resulting pseudotime was low in NPCs, high in neurons, and intermediate in other cell types, as expected (Fig. 4B). The mesenchymal lineage represents a distinct branch from IPC, indicating a divergent fate rather than a transitional state between NPCs and neurons. The homozygous-specific cluster consisted mostly of Mes-KO and NPC (Fig. 4C, circled). *RAI1*-homozygous cells showed higher global pseudotime (Fig. 4C&4D). The higher pseudotime of *RAI1*-homozygous cells was observed in NPCs, mesenchymal cells, and IPC of day 20, but not in neurons and astrocytes of day 60 (Fig. 4E). *RAI1*-heterozygous cells showed similar pseudotime distribution with WT. These results indicate that advanced differentiation states in *RAI1*-homozygous mutants are not cell-type-specific but rather a general feature at day 20.

**Figure 4.**
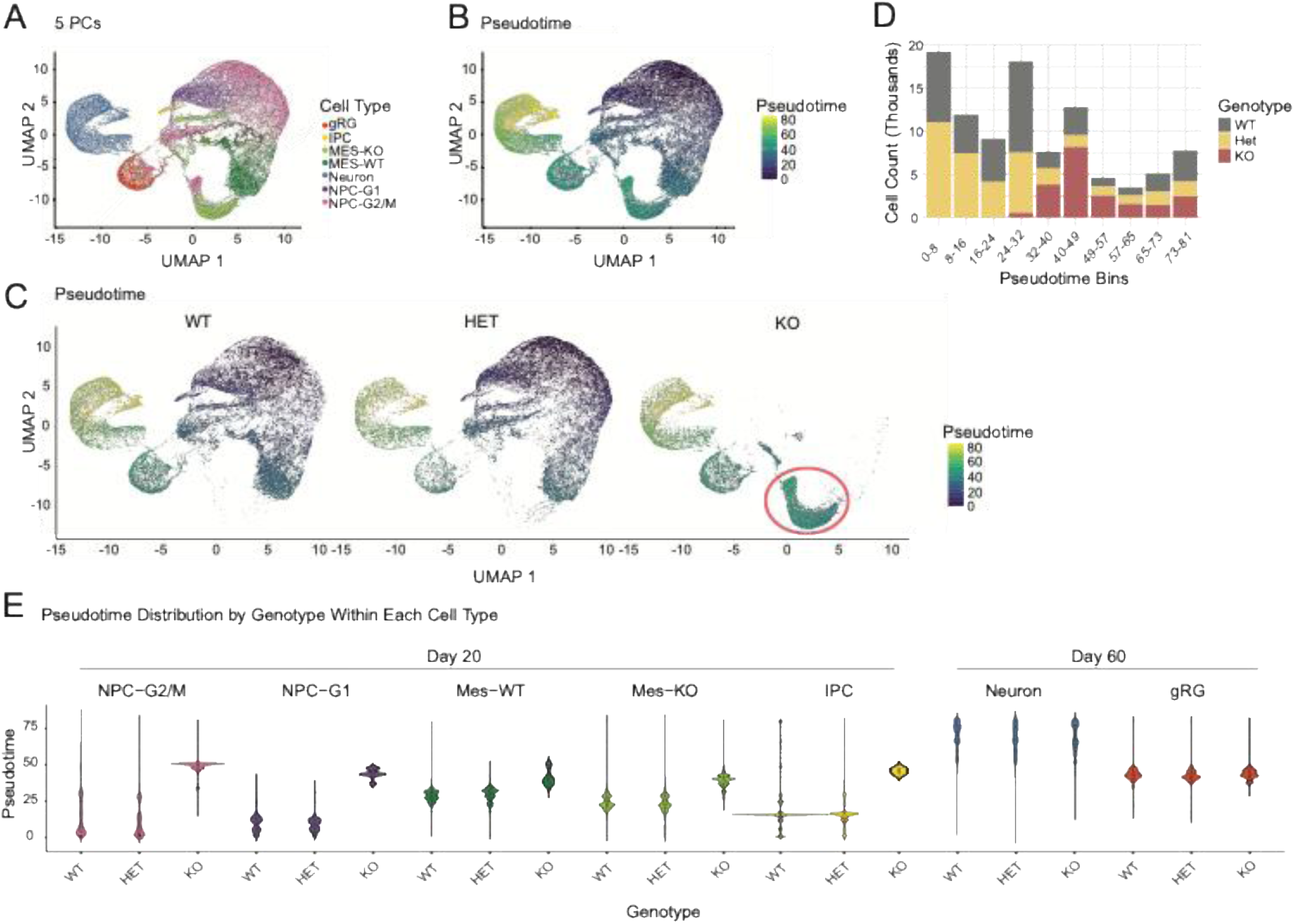
Pseudotime analysis of scRNA-seq data. **(A)** A single-cell transcriptome trajectory constructed with Monocle 3 (26). Cell types assigned with Seurat, as described earlier, are overlaid. **(B)** Pseudotime distribution: Randomly chosen 10 WT NPC-G2/M cells (UMAP1 > 5 and UMAP2 > 5) were set as the roots. **(C)** Pseudotime UMAP plots split by genotypes. **(D)** Cell numbers across pseudotime bins. **(E)** Pseudotime across cell types.

### Day 20: Transcriptional signatures of ectoderm-to-mesoderm derailment

To gain insights into potential cell biological consequences of *RAI1* loss, we investigated the genes dysregulated in *RAI1*-KO cells across cell types using pseudobulk differential gene expression analysis (27). The number of differentially expressed genes (DEGs, Padj <0.01, |Log2FC| > 1, list: Table S2&S3) was greater in homozygous mutants compared to heterozygous across cell types, as expected (Fig. 5A). Among the cell types, NPCs, Mes, and IPCs exhibited greater numbers of DEGs compared to those within gRG and neurons, consistent with bulk RNA-seq data (Fig. 2) and pseudotime analysis (Fig. 3), collectively demonstrating that the impact of *RAI1* loss is greater in the neuroepithelial phase (day 20) than the post-mitotic/maturation phase (day 60).

**Figure 5:**
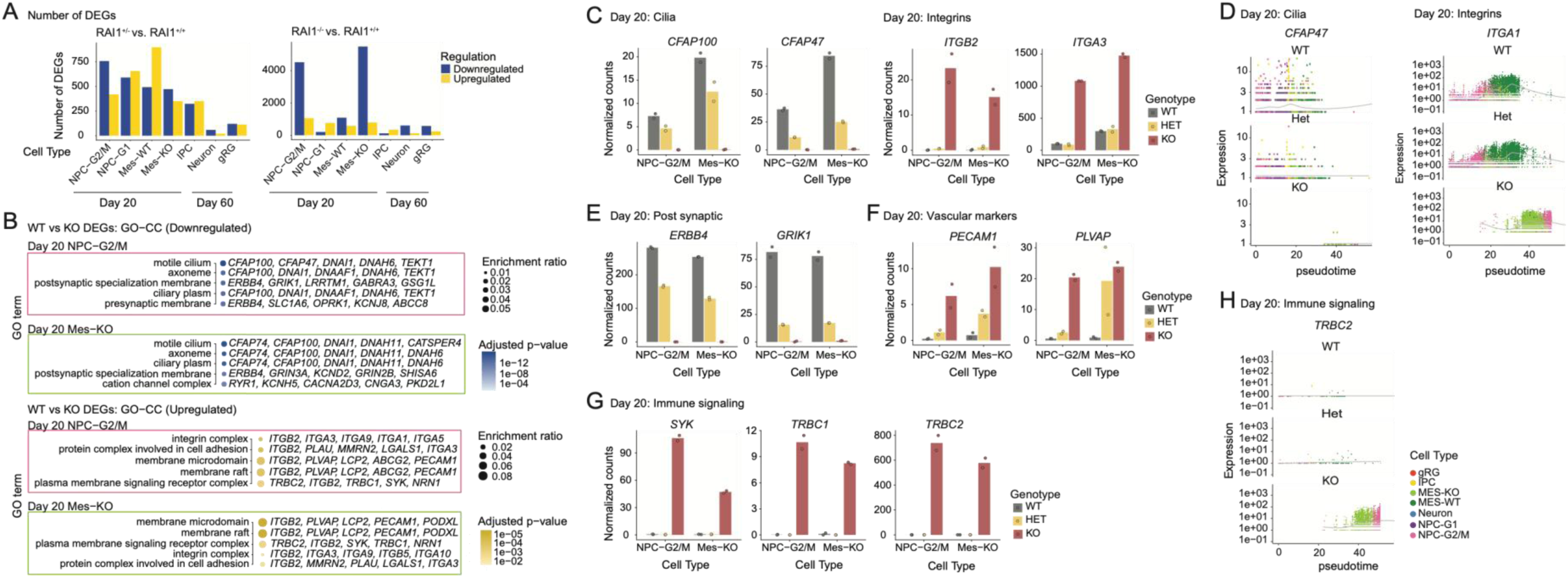
Genes dysregulated in *RAI1*-deficient cell types on day 20. **(A)** The number of differentially expressed genes in each cell subtype (Padj < 0.01, Log2FC > 1 or < −1). **(B)** GO Cellular Component enrichment for *RAI1*-homozygous DEGs. The top five GO terms are presented with the genes that contributed most to the enrichment. **(C)** Expression of the top two cilia and the top two integrin genes is plotted across genotypes. **(D)** Expression of *CFAP47* and *ITGA1* across the pseudotime. The color of dots represents the cell types denoted in Figure 4A. **(E-G)** Expression of the top two synaptic, vascular, and immune signaling genes across the gene types. **(H)** Expression of the T-cell receptor gene *TRBC2* along the pseudotime across the genotypes. Mean values are shown in histograms, with individual data points from biological duplicates. The day 20 data are plotted, and the expression of additional representative genes is presented in Figure S5.

With the day 20 dataset, gene ontology analysis shows that *RAI1*-homozygous NPCs and mesenchymal cells exhibit downregulation of genes encoding primary cilia components (e.g., *CFAP100* and *CFAP47*), accompanied by upregulation of integrins (e.g., *ITGB2*, *ITGA3*) (Fig. 5C, Fig. S5A). As primary cilia form at the apical membrane of NPCs while integrins define the basal membrane (28), the pattern is suggestive of impaired cell polarity. Plotting cilia and integrin gene expression across pseudotime revealed that no *RAI1*-homozygous cell types expressed cilia genes. Meanwhile, integrin genes play important roles in the homing of cells that underwent epithelial-mesenchymal transition (EMT) (28). Together, *RAI1* homozygous deletion leads to transcriptional signatures of enhanced EMT processes.

The down-regulation of synaptic factors (e.g., *ERBB4* and *GRIK1*, Fig. 5B&5E, Fig. S5A) initially appeared discordant with the advanced pseudotime of *RAI1*-homozygous mutant NPCs (Fig. 3). A close examination of the upregulated membrane genes revealed that they encode vascular endothelial markers (e.g., *PECAM1*, *PLVAP*) and immune cell proteins (e.g., *SYK*, *TRBC1*), which should not be expressed in NPCs or their derivatives (Fig. 5F&5G, Fig. S5A). The ectopic gene expression progressed along pseudotime (Fig. 5H), suggesting that the mesodermal genes drive advanced pseudotime at day 20. The initial separation of Mes-WT and Mes-KO clusters also involves the loss of neuronal genes and the ectopic immune gene expression in Mes-KO (Fig S5B). Thus, at day 20, *RAI1* -homozygous NPC and mesenchymal cells exhibited a sign of lineage derailment from the ectoderm to the mesoderm.

In *RAI1*-heterozygous mutant cells, the ectopic expression of integrin genes and most immune cell genes was not observed (Fig. 5C, Fig. S5A). However, heterozygous mutant cells exhibited reduced neural gene expression and ectopic expression of vascular and some immune genes (e.g., *BST2*, Fig. S5A), suggesting a mesodermal shift in heterozygous cells to some extent. Gene ontology analysis of *RAI1*-heterozygous mutant cell types identified similar groups of deregulated genes, while additional ectopic expression of myogenic genes (e.g., *ACTA1*, *ACTC1*) and gastrointestinal tract genes (*CLDN1*, *CLDN4*) was observed (Fig. S5C&D). These mesoderm-like (some endoderm-like in heterozygous) signatures were common across cell types. Together, during early neural differentiation, the greatest impact of *RAI1* loss manifests as ectopic mesodermal or endodermal gene expression, rather than dysregulation of neural cell type-specific gene expression programs.

### Day 60: Transcriptional signatures for enhanced neuronal maturation

We next examined gene expression changes in gRG and neurons at day 60. Within gRG, downregulated genes are enriched for pathways involving the neural crest-derived cells (Fig. 6A). For example, *PMEL*, *TYR*, *ABCC9*, *LAMA2*, and *NID1*, which contributed to the downregulated pathways, are normally expressed in the choroid plexus, pericytes, and meninges, all of which have contributions from neural crest (29–32). In contrast, the upregulated pathways in *RAI1*-homozygous mutants were primarily associated with neuronal maturation in both gRGs and neurons (Fig. 6A). These include synaptic genes *NRGN, ITGA8, SHISA9, GRM5,* and *GRIA4* (Fig. 6B) as well as mitochondrial genes, such as *MT-ND5, MT-ND2, MT-ATP6, MT-ND1, MT-CO2*, many of which encode subunits of Complex I of the mitochondrial electron transport chain (Fig. 6C). A metabolic change from aerobic glycolysis to oxidative phosphorylation (Oxphos) is a hallmark of neural maturation (33,34). Higher expression of mitochondrial genes can indicate oxidative stress; however, we did not detect significant increases in reactive oxygen species scavengers, mitophagy genes, or mitochondrial unfolded protein response genes (Fig. S6A-C).

**Figure 6:**
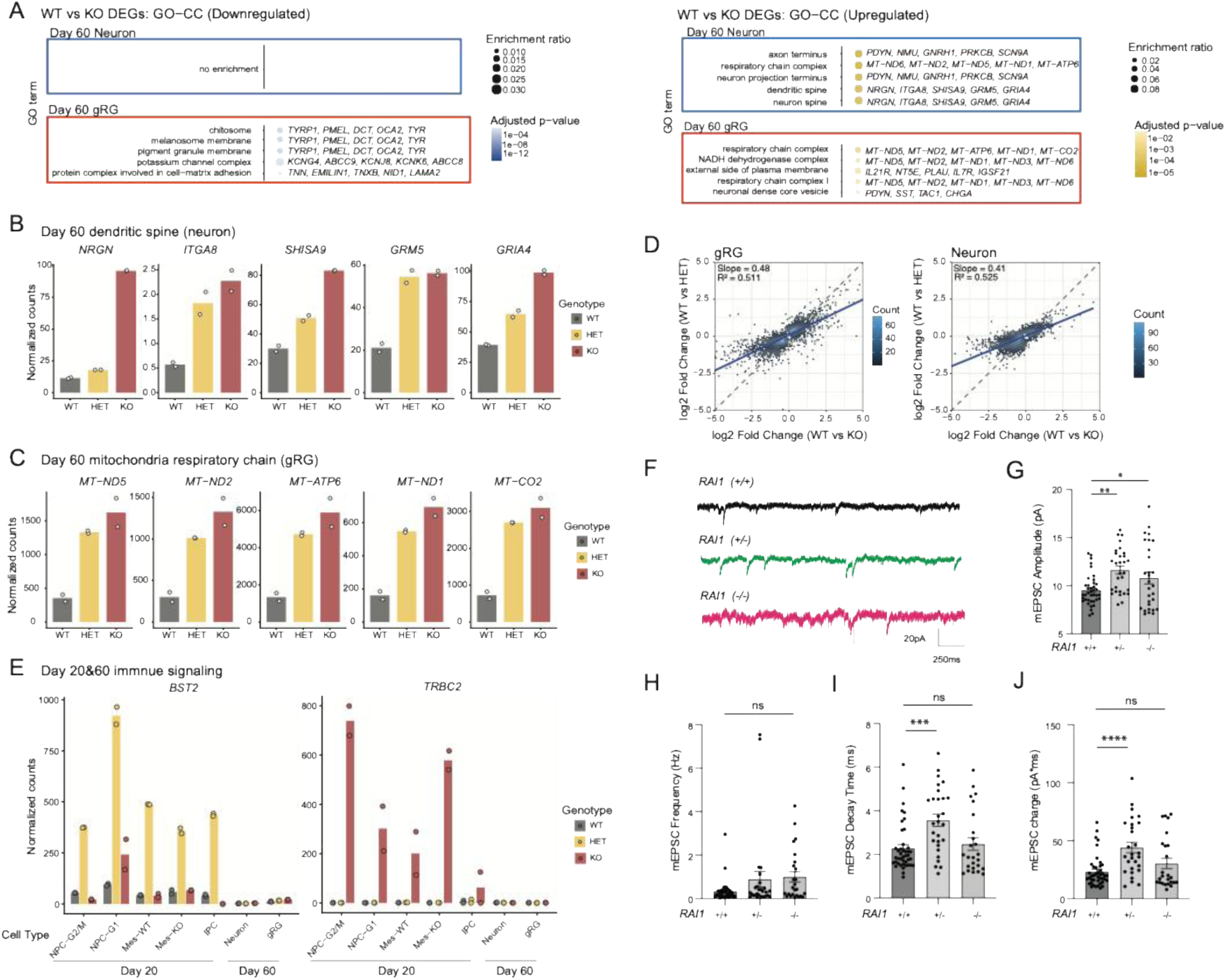
Genes dysregulated in *RAI1*-deficient cell types on day 60. **(A)** GO Cellular Component enrichment for *RAI1*-homozygous DEGs. The top five GO terms are presented with the genes that contributed most to the enrichment. **(B&C)** Expression of the top 5 dendritic spine genes in neurons and respiratory chain complex genes in gRG across genotypes. **(D)** Comparison of the magnitude of changes of all DEGs, identified in *RAI1*-heterozygous and homozygous mutants, between the two genotypes. **(E)** Immune signaling genes were derepressed only on day 20 but not on day 60. Additional genes shown in Fig. S6. Mean values are shown in histograms, with individual data points from biological duplicates. **(F-J)** Analysis of mEPSCs recorded from *RAI1*+/+ (n = 42), *RAI1*+/-(n = 28) and *RAI1*-/-(n = 25) neurons. Mean ± s.e.m. are shown in bar graph, with individual data points. Statistical analysis were performed using one-way ANOVA followed by Tukey’s post hoc tests. Significance is indicated as *p < 0.05, **p < 0.01, ***p < 0.001, ****p < 0.0001; ns p ≥ 0.05. **(F)** Representative mEPSC traces. **(G)** mEPSC amplitude. **(H)** mEPSC Frequency. **(I)** mEPSC decay time. **(J)** mEPSC charge.

Meanwhile, the expression of nuclear-encoded Oxphos factors tended to be increased in *RAI1*-mutants (Fig. S6D). The upregulation of mitochondrial genes and neuronal maturation genes was clearly greater in homozygous mutants compared to heterozygous (Fig. 6B), and this *RAI1*-dose dependency extended to all DEGs (Fig. 6D). Mesodermal genes, including immune signaling genes, de-repressed in day 20 mutant cells, were no longer expressed at day 60 (Fig. 6E&S6E). Thus, in contrast to the pro-mesoderm signature at day 20, *RAI1*-deficient cells at day 60 showed a transcriptional signature of pro-neuronal, accelerated neuronal maturation.

To examine whether accelerated neurodevelopmental gene expression programs are associated with altered functional maturation, we measured excitatory synaptic function at day 60. We used whole-cell patch-clamp recordings of miniature excitatory postsynaptic currents (mEPSCs) to measure synaptic currents at a time point where excitatory synapses are initially forming. Cells with a pyramidal-like neuron morphology were voltage-clamped at -70 mV in the presence of a GABA-A receptor antagonist (10 μM bicuculline) and the sodium channel blocker tetrodotoxin (1 uM) to isolate AMPA receptor-mediated events. Consistent with accelerated synaptic maturation, we found a robust increase in mEPSC amplitude in both heterozygous and homozygous *RAI1* mutant neurons relative to controls (Fig. 6F&6G). Similarly, mEPSC frequency showed a trend toward an increase in both heterozygous (p = 0.1617) and homozygous (p = 0.0871) neurons (Fig. 6H), suggesting that, in addition to being stronger, excitatory synapses were also more numerous with *RAI1* deficiency. Unlike the *RAI1* dose-dependent increase in synaptic gene expression, the mEPSC phenotype was not stronger in *RAI1*-homozygous mutants relative to heterozygous mutants, which could reflect a ceiling effect.

Together, these results suggest that accelerated neurodevelopmental gene expression is accompanied by faster maturation of excitatory synapses in *RAI1* mutant neurons.

### *Rai1*-deficient murine cells do not show an accelerated differentiation signature

Having established *RAI1*’s role in slowing the pace of human cortical cell differentiation, we sought to test whether this function is conserved in mice. We first examined the kinetics of neuronal differentiation gene expression in ESC-derived NPCs/neurons with homozygous *Rai1* deletion and WT control (35). This RNA-seq dataset includes two time points—day 8 (NPCs) and day 12 (early post-mitotic neurons). To place the available time series within a developmental transcriptome trajectory, we generated mRNA-seq data from undifferentiated WT mouse ESCs and day 7 cortical neuron cultures derived from E16.5 embryos (referred to as day 23), representing more mature neurons. PCA analysis of the integrated datasets showed no sign of either delay or acceleration in differentiation (Fig. 7A). The orthologues of our cluster 3 synaptic genes (Fig. 2H), which represented accelerated neuronal maturation in *RAI1*-homozy cells, did not show any changes in *Rai1*-KO mouse NPC/neurons (Fig. 7B). Differential gene expression analysis comparing *Rai1*-KO and WT mouse cells indicated that the gene dysregulation primarily occurred at day 12 (Fig. 7C). Downregulated genes are enriched with protein folding stress pathways (e.g. *Hspa1a/b*), and upregulated genes are enriched with lipid sensing pathways (e.g. *Lsr, Apoe*, and *Selenos*), with no obvious link to differentiation states (Fig. 7D).

**Figure 7:**
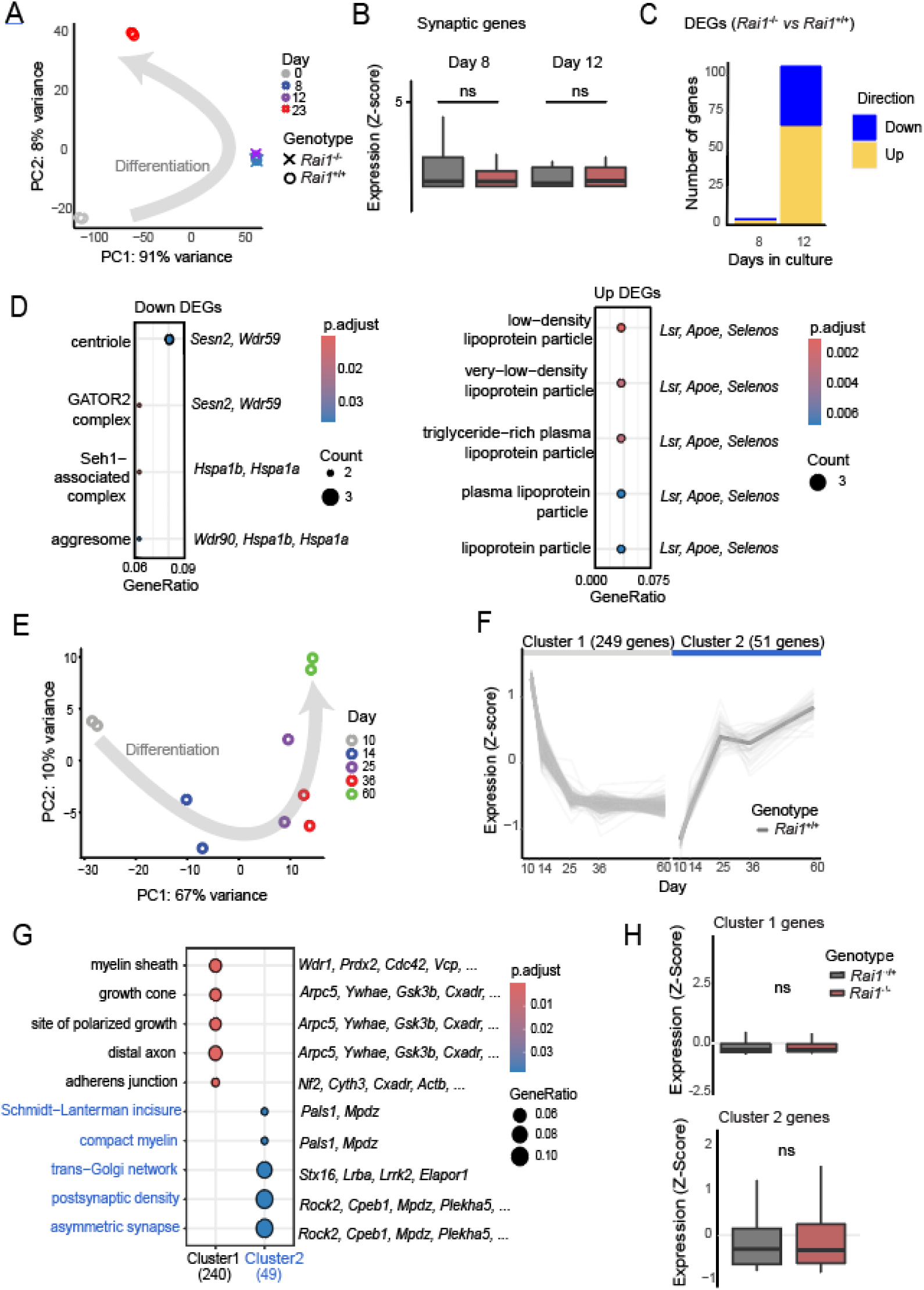
*Rai1*-KO in mice does not alter developmental tempo *in vitro* nor *in vivo*. **(A)** Developmental trajectory of Kasper *et al* data nested in ESC and DIV7 neuron data within PC space. **(B)** Gene expression of mouse orthologs of the Cluster 3 human genes that showed acceleration in *RAI1*-KO in Fig.2H. **(C)** Differentially expressed gene (DEG) numbers in Kasper *et al* data. **(D)** GO cellular component analysis of upregulated and downregulated DEGs from part C. **(E)** Developmental trajectory of wildtype postnatal mouse cortex in PC space. Data was obtained from ENCODE database. **(F)** Gene expression in clusters identified from top 300 loading PC1 genes from part E. **(G)** GO cellular component analysis of gene clusters. **(H)** No significant difference in either cluster gene expression in P21 *Rai1-*cKO (bulk RNAseq data from Huang *et al*)

Next, we examined *in vivo* mRNA-seq data from the P21 cortex of brain-targeted *Rai1*-cKO with Nestin-Cre (18). This dataset represents a single time point, and the transcriptome did not align closely with the longitudinal ENCODE mRNA-seq data from mouse cortices at P10, P14, P25, P36, and P60 in PCA space, likely due to batch effects (not shown). We reasoned that we could first identify neuronal maturation genes using WT ENCODE data, then test whether these genes change in *Rai1*-KO brain and cells at P21. As proof of principle, we first applied this strategy to our data and found that it enabled inference of differentiation states in mutant cells (Fig. S7; see Methods). Principal component 1 of the ENCODE dataset explained 67% of the variance in the transcriptome across the five developmental time points (Fig. 7E). K-means clustering of the top 300 PC1 high-loading genes identified two clusters: cluster 1 represents downregulated genes, and cluster 2 represents upregulated genes during neuronal maturation (Fig. 7F). Cluster 1 was enriched for early developmental components, such as growth cones, while Cluster 2 was enriched for later maturation structures, such as postsynaptic density (Fig. 7G). However, neither cluster showed differential expression in the cortex of the P21 *Rai1*-cKO mouse (Fig. 7H). Collectively, published mouse *RAI1*-mutant mRNA-seq data, both *in vitro* and *in vivo,* show no evidence of altered temporal dynamics of neurodevelopmental gene expression.

### Loss of *RAI1* accelerates gene expression program of human iNeuron differentiation

We then sought to determine the impact of *RAI1* loss in iNeurons in which the neurogenic transcription factor NGN2 is expressed via the TetON inducible promoter (36,37) (Fig. 8A, see Method). NGN2 expression has been shown to significantly shorten the time for neuronal differentiation and maturation. Initially, we expected that the forced NGN2-driven differentiation would mask the accelerated differentiation phenotype of *RAI1*-deficient neurons. Contrary to our prediction, bulk RNA-seq of *RAI1*-deficient iNeurons at three time points (day 0, 3, and 7) showed accelerated differentiation even more pronounced than that following the dual SMADi (Fig. 8B&C). Both heterozygous and homozygous mutants showed advanced differentiation transcriptome states relative to controls, with a larger effect in homozygous mutants. The *RAI1* dose-dependent effect is corroborated by the higher number of DEGs (Padj < 0.05, list: Table S4) in homozygous than in heterozygous mutants at day 3 and day 7 (Fig. 8D&E). Hierarchical clustering of DEGs yielded four clusters. Of these, cluster 2 showed accelerated induction in *RAI1*-mutant cells and was enriched with synaptic genes (Fig. 8F&G), supporting the faster differentiation. Cluster 3, downregulated in mutants, represents replication machinery, suggesting earlier cell cycle exit.

**Figure 8.**
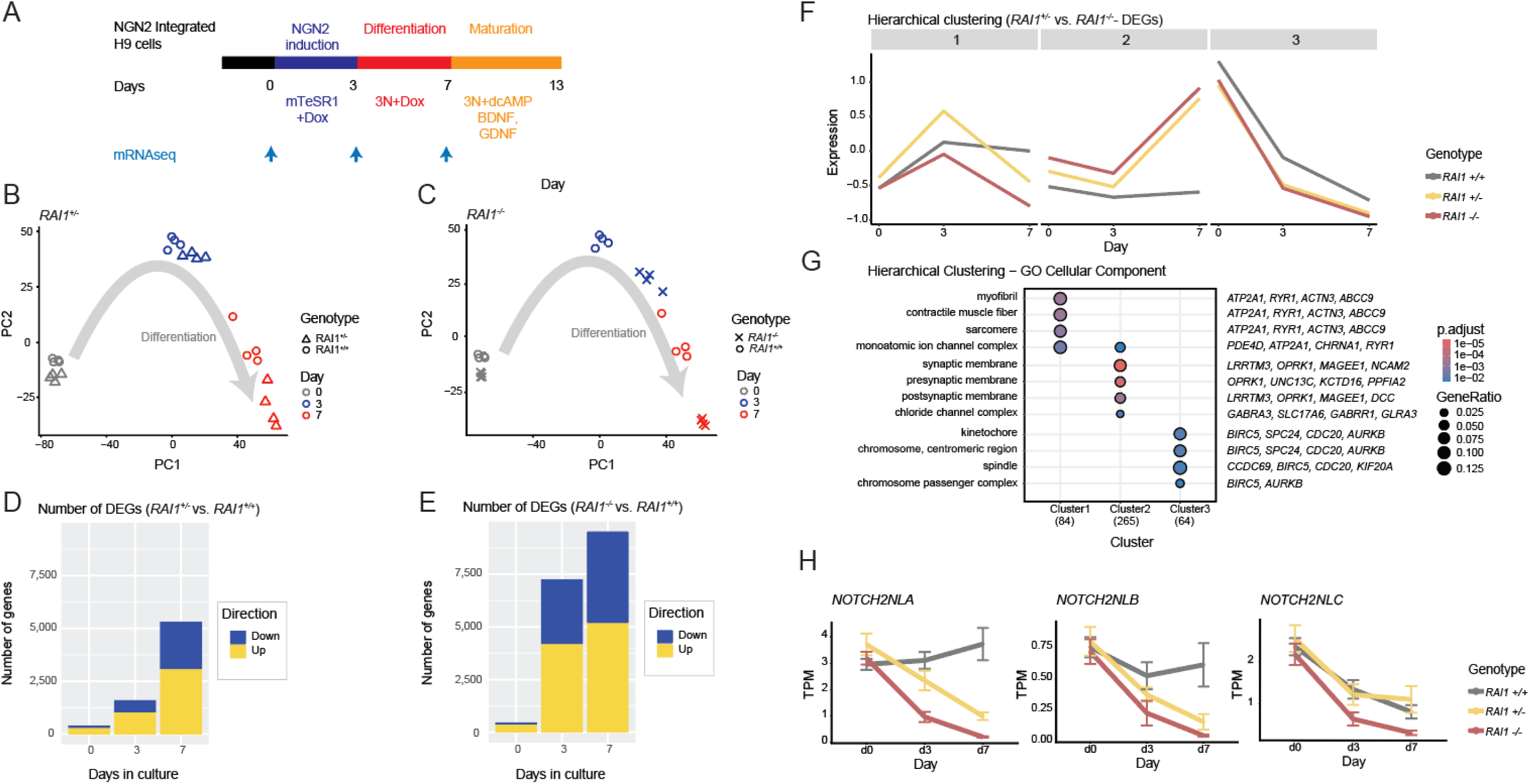
Accelerated differentiation of RAI1-deficient iNeurons. **(A)** Experimental scheme. **(B&C)** PCA analysis of the transcriptome data (C: WT & *RAI1*+/-, D: WT & *RAI1*-/-). **(D&E)** DEG numbers (E: WT vs. *RAI1*+/-, F: WT vs. *RAI1*-/-). **(F)** Hierarchical clustering of DEGs followed by Molecular Function GO analysis. **(G)** GO Cellular Component analysis of DEG cluster genes. The parentheses indicate the number of genes. **(H)** The expression of human genes is implicated in the slower tempo of brain development.

To gain insight into potential mechanisms underlying accelerated neurodevelopmental gene expression patterns, we systematically examined 19 human- and primate-specific genes and their regulators implicated in the slower tempo of human brain development (1,38) (Fig. S8A). Of these genes, *NOTCH2NL* family genes were significantly down-regulated in *RAI1*-deficient iNeurons (Fig. 8H). *NOTCH2NL* family genes suppress transition from neural progenitor to neurons, thereby expanding and increasing neural numbers in the human brain (6,39); therefore, these are potential candidates as mediators of *RAI1*’s roles in slowing the pace of neurodevelopment. 12 genes were covered by single-cell RNA-seq probes, and their expression shows mostly downregulation in *RAI1*-deficient cell types, with a greater effect in the NPC stage, but some, such as *ZEB2*, are downregulated in mature neurons (Fig. S8B).

Together, *RAI1*’s dose-dependent function suppresses the gene expression program of human neuronal differentiation and maturation. The iNeuron results isolate the roles of *RAI1* as a brake on the neurodevelopmental gene expression program at the NPC stage, which is confounded by the ectopic expression of mesoderm/mesenchymal genes in the dual SMADi system (Fig. 5). Additionally, a more pronounced effect in iNeurons suggests a functional interplay between NGN2 and *RAI1*.

## Discussion

Our work provides multiple lines of evidence for *RAI1*’s role in slowing the tempo of human neurodevelopmental gene expression programs. *RAI1* deficiency accelerates developmental transcriptome changes across three sequencing studies: bulk and single-cell RNA-seq of dual SMADi cortical differentiation and iNeurons. Notably, this function appears to be human-specific, as mouse *Rai1*-deficient cells and tissues did not exhibit comparable acceleration of differentiation signatures. Additionally, single-cell RNA-seq analysis revealed that *RAI1* suppresses mesodermal and endodermal gene expression at early stages. These findings reveal *RAI1* as a key regulator of the extended developmental timeline characteristic of human brain development and as a guardian of ectodermal identity.

Human neurodevelopment progresses through at least two stages: NPC expansion and neuronal maturation. For mouse NPC expansion, chromatin factors such as HMGN1 (40), ZNF335 (41), GLI3 (42), and PRDM16 (43) have been shown to protect NPCs from premature differentiation, but their roles in human NPCs have not been experimentally tested. For neuronal maturation, a recent study uncovered a dozen “barrier” chromatin regulators that slow down the process (8). In the present study, the data clearly demonstrate the roles of *RAI1* in the slow maturation program because both *RAI1*-heterozygous and homozygous mutants display faster expression of synaptic genes (Fig. 2, 5 &6), which is corroborated by electrophysiological evidence (Fig. 6). The roles of *RAI1* in the neurodevelopmental tempo at NPC is less clear because, in the NPC stage, the ectopic expression of mesodermal genes confounds the assessment of neurodevelopmental tempo (Fig. 4&5). However, the clear transcriptomic acceleration of the early differentiation phase in *RAI1*-mutant iNeurons (day 3, when cells exit the cell cycle) supports the model in which *RAI1* extends both the NPC expansion and maturation phases.

How does *RAI1* prolong the neurodevelopmental gene expression program? A key distinction between *RAI1* and previously identified barrier chromatin factors is their temporal expression patterns: whereas barrier chromatin factors are downregulated during neuronal maturation (8), *RAI1* expression increases (Fig. 2). Barrier transcription factors typically repress pro-maturation genes, and this repression is subsequently relieved to permit maturation. The progressive upregulation of *RAI1* during neural development (Figure 2B) suggests that *RAI1* functions as a rheostat that modulates the pace of neuronal maturation. In this capacity, *RAI1* may track increases in gene expression and constrain them to prevent premature or excessive induction. A second notable finding is that *RAI1* loss exerts a more pronounced effect on neurodevelopmental tempo in the iNeuron system (Fig. 8) than in the dual-SMAD inhibition system (Fig. 2&4). These results suggest mechanistic interactions between RAI1 and NGN2. For example, RAI1 may attenuate NGN2-mediated transcriptional activation, thereby preventing excessive transcriptional output. Future investigations should elucidate the molecular mechanisms by which RAI1 prolongs human neurodevelopmental programs.

The present study highlights mesenchymal/neural crest cells as an important cell type for investigating *RAI1*’s role in Smith-Magenis syndrome. Mesenchymal cells appeared only transiently in our culture on day 20 (Fig 3), likely due to culture conditions optimized for neurons. This cell population provided an opportunity to assess *RAI1*’s role in the developmental tempo of non-neuronal cells. Neural crest is a multipotent, migratory cell population that departs from the neural plate via epithelial-mesenchymal transition and gives rise to the peripheral nervous system, melanocytes, and craniofacial bones and skin. In the central nervous system (CNS), the neural crest gives rise to the choroid plexus, pericytes, and meningeal fibroblasts that support collagen-rich meningeal membranes and to pericytes that wrap around vascular capillaries (44). Notably, *RAI1* loss appears to have a stronger impact on neural crest, as the *RAI1* homozygous culture showed a distinct cluster with a more advanced pseudotime, driven by a mesodermal gene expression signature at day 20 (Figs 3-5). In addition, at day 60, genes involved in CNS neural crest cell development are downregulated in *RAI1*-mutants (Fig. 6). Thus, investigating the roles of *RAI1* in underappreciated neural crest-derived cell types, such as meningeal fibroblasts and pericytes, will likely provide important insights into SMS pathophysiology.

Given that mechanisms of chromatin regulation are highly conserved across taxa, the fundamental assumption can be that their roles in neurodevelopment are also conserved. Indeed, key SMS features have been recapitulated in mouse models, including obesity, circadian abnormalities, and decreased social behavior (15,45,46). However, our transcriptome analysis of available data from *Rai1*-deficient mice did not reveal conserved roles for *RAI1* in developmental gene expression. This differential impact of *RAI1* loss on mouse and human has been observed at both NPC and neuronal maturation stages. Some SMS individuals exhibit ventriculomegaly, and NPCs derived from SMS iPSCs with 17p11.2 deletion showed enlarged ventricles accompanied by accelerated cell-cycle exit, which likely underlie ventriculomegaly (19), both of which can be attributed to *RAI1* deficiency based on our data. *Rai1*-deficient mice lack such an NPC phenotype; they have a normal brain structure (15,45,46). At the synaptic level, SMS iPSC-derived cortical neurons exhibit increased excitatory synapse formation (19) (Fig. 6), while pyramidal neurons in the *Rai1*-KO mouse prefrontal cortex showed reduced dendritic spine density (46), with the caveat that these are not exactly the same measurements.

The impact of *RAI1* loss on neural crest appears to differ between species as well. Human SMS individuals frequently exhibit craniofacial (>75%) and dental (>90%) abnormalities attributable to neural crest deficiency (13). Craniofacial abnormalities in *Rai1*-heterozygous mice have a low penetrance; larger deletions surrounding *Rai1* increased the penetrance, implicating a contribution of the *Rai1* regulatory region or other genes in the SMS interval (47). In contrast, *Rai1* knockdown in Xenopus was sufficient to cause midface hypoplasia and mouth deformity reminiscent of human SMS, along with neural crest reduction and migration deficiency (48). Interestingly, DNA sequence changes in canine *RAI1* regulatory regions near its promoter have been linked to the domestication of dogs, during which many neural crest-associated morphological changes occurred (e.g., from a uniform pointy snout to diverse features across breeds) (49). These observations, together with the present data, imply that *RAI1*’s roles diverged relatively recently for neural crest-related developmental processes.

### Limitation

Our study is primarily gene expression analyses; therefore, it can provide limited insights into the cellular impact of RAI1 deficiency except for the mEPSC measurements. Instead, the presented results provide testable hypotheses regarding the cell-biological consequences, for example, the lack of primary cilia in mutant NPC and the functional deficits of neural crest cells. In addition, although transient, the misexpression of mesodermal genes could entail irreversible damage to cellular function. Future studies should test these predictions and hypotheses.

## Experimental procedures

### Cell Culture and *RAI1* knockout cell line generation

hESCs H9 cells were cultured on Geltrex-coated plates (Gibco A1413302) and fed with TeSR-E8 medium. 10 μM ROCK inhibitor Y27632 (Cayman 100005583) was added during cell passage or thawing. *RAI1* knockout cell lines were generated using the CRISPR/Cas9 system. Briefly, we designed high-scoring gRNAs based on Optimized CRISPR design (50).sgRNA: TGTGGTCAGTGATCCCTGCGGGG, which targets exon 3, was selected for the following experiment. The synthesized oligos of gRNAs were annealed and cloned into the pX459 vector carrying Cas9 and a gRNA expression cassette (51). The recombinant plasmid was then transfected into H9 cells using Lipofectamine (Invitrogen). Single clones were isolated after puromycin selection, and the genotype was confirmed by Illumina sequencing of PCR products from genomic DNA. Each line was assessed for chromosomal abnormalities by single-nucleotide polymorphism (SNP) chip microarray (Infinium Core Exome-24 BeadChip, Illumina, San Diego, California, USA) and analyzed by Genome Studio 2.0 (Illumina), using hg19 as the reference genome.

### Cortical neuron differentiation

Cortical neurons were differentiated following a modified dual-SMAD inhibition protocol (23). Briefly, 1 million cells were plated in each well with TeSR1/E8 medium to initiate the experiment (day 1). From day 0–11, cells were cultured in 3N media without vitamin A supplemented with 1 µM dorsomorphin (Sigma P5499) and 10 µM SB431542 (Cayman 13031). On day 12, cells were passaged using dispase and plated at 1:3 onto Geltrex-coated 6-well plates in 3N medium containing dorsomorphin and SB431542. The 3N medium was then replaced daily. When rosettes appeared, 20 ng/mL FGF2 was added for 4 days. On day 20, cells were passaged again using dispase, and 3N medium was replaced every other day. On day 27, cells were passaged using Accutase and cultured in 3N medium with vitamin A, with medium changes every other day. On day 34, cells were passaged again using Accutase, and 0.5 million cells were plated in 3N medium with vitamin A on PLO/laminin-coated 6-well plates. During the first week post-plating, the medium was supplemented with 0.2 µM Compound E (EMD BioSciences 565790), 20 µg/mL GDNF (PeproTech 450-10), 20 µg/mL BDNF (PeproTech 450-02), and 0.4 mM dbcAMP (Sigma D0627-1). During the second week, the medium contained 20 µg/mL GDNF, 20 µg/mL BDNF, and 0.4 mM dbcAMP. From the third week onward, only 20 µg/mL GDNF and 20 µg/mL BDNF were added to the medium. Half of the medium volume was changed every 3–4 days. Every 10 days, a full-volume medium change was performed, and 1 µg/mL laminin was added.

### iNeuron generation and differentiation

We generated iNeuron hESC lines carrying doxycycline-inducible NGN2 transgene as previously described (37). H9 hESC lines (WT and *RAI1* mutants) were typically transfected with 2.5 μg of total DNA using 7.5 μL of Lipofectamine Stem Reagent (Thermofisher STEM00008). To generate new glutamatergic iNeuron lines, we used 0.75 μg of pCAGGS-PBase, 0.25 μg of PB-CAG-puro, 0.75 μg of PB-CAG-TetOn-IRES-mCherry, and 0.75 μg of PB-TRE-hNGN2. The transfection complex was prepared in OptiMEM according to the transfection reagent protocol. Three days after transfection, cells were passed to a 10 cm dish. After 6 days, 0.85 μg/mL puromycin or 200 μg/mL G418 was added to the medium. Clones were picked once the cells grew to colonies of 100–500 cells.

On day -1, iPSCs were dissociated with Accutase and each line plated in 3 wells of a Geltrex-coated 6-well plate at a density of 0.5 - 1 × 10^6^ cells per well in mTeSR Plus (StemCell Technologies) with 10 μM ROCK inhibitor Y27632. Cell lines were plated at the same density for each differentiation. The following day (day 0), media was changed to mTeSR Plus with 1 μg/mL doxycycline (Sigma D9891). Media was changed to fresh mTeSR with 1 μg/mL doxycycline on day 1. On day 2, media was changed to Neurobasal-A supplemented with 1x Glutamax, 1x B27 supplement, 20 ng/ml GDNF, 20 ng/ml NT3, 1x CultureOne, and 1 μg/mL doxycycline. Starting on day 3, media was switched to Neurobasal-A with 1x Glutamax, 1x B27 supplement, 20 ng/ml GDNF, 20 ng/ml NT3, and 1x CultureOne and changed every other day. RNA was collected from one well for each line on days 0, 3, and 7 for bulk RNA sequencing.

### Bulk mRNA-seq and data analysis

Bulk mRNA-seq was performed as 2×10⁵ human cells were collected for each sample. RNA was extracted using TRIzol reagent (Ambion) and purified using the RNeasy Mini kit (QIAGEN) according to the manufacturer’s protocol. Similarly, mouse ESCs and cortical neuron mRNAseq data were generated using cells cultured as described previously (21,52). Total RNA from each sample was used to prepare Illumina sequencing libraries with oligo-dT priming. The sequencing data are processed in our previously described pipeline (53). For iNeuron, bulk RNA-seq Libraries were prepared using NEBNext rRNA Depletion Kit (Human/Mouse/Rat) v2 (NEB #E7400X), and NEBNext UltraExpress RNA library prep kits (NEB #E3330L), with the following adapter sequences: R1: AGATCGGAAGAGCACACGTCTGAACTCCAGTCA and R2: AGATCGGAAGAGCGTCGTGTAGGGAAAGAGTGT. Sequencing was performed on an Illumina NovaSeqXPlus platform using 151bp paired-end reads according to the manufacturer’s protocol. De-multiplexed FASTQ files were generated using BCL Convert Conversion Software v4.0 (Illumina). The bioinformatics workflow was managed using Snakemake (54). Raw reads were trimmed using Cutadapt v4.8 (55) and quality-assessed with FastQC v0.11.8. Trimmed reads were aligned to the human reference genome GRCh38 (ENSEMBL 109) using STAR v2.7.8a (56) following ENCODE standards for RNA-seq. Gene-level count estimates were generated using RSEM v1.3.3 (57). Quality control metrics from multiple pipeline steps were aggregated using multiQC v1.20 (58). The sequencing quality matrix is presented in Table S5.

### Single-cell RNA sequencing and data analysis

Single-cell RNA sequencing libraries were prepared with Chromium Fixed RNA Profiling Platform (Flex, 10x genomics). For each sample, 3×10⁵ cells were fixed using the NEXT GEM Cell Fixed RNA Sample Preparation Kit (10x Genomics, 1000414). Cells were washed with ice-cold PBS + 0.04% BSA. Then, 1 ml Fixation Buffer was added to the washed cell pellet and mixed by pipetting, followed by overnight incubation at 4°C. Samples were centrifuged at 850 rcf for 5 minutes at room temperature, and 1 ml ice-cold Quenching Buffer was added and mixed by pipetting. Next, 100 µl pre-warmed Enhancer was added to the 1 ml fixed sample in Quenching Buffer, followed by 275 µl 50% glycerol for storage. Prepared samples were submitted to the University of Michigan Advanced Genomics Core for library preparation and sequencing. The pooled library was sequenced for 28x91 bp according to the manufacturer’s protocol (Illumina NovaSeq 6000). BCL Convert Conversion Software v4.0 (Illumina) was used to generate de-multiplexed Fastq files. The cells with less than 20% mitochondrial reads were used for further analysis. The sequencing quality matrices are reported in Table S6.

### Analysis of mouse RNAseq datasets

Bulk RNA-seq data from the female mouse cortex were obtained from the Encyclopedia of DNA Elements (ENCODE) portal (59–61) using the following accession identifiers-Day 10: ENCFF557BVU, ENCFF429GAO; Day 14: ENCFF585AWT, ENCFF455XPL; Day 25: ENCFF759EMR, ENCFF476AFO; Day 36: ENCFF031XSD, ENCFF068HYN; Day 60: ENCFF806DOB, ENCFF404SAA. The raw data were deposited to ENCODE by the Barbara Wold laboratory, Caltech. Bulk RNA-seq data from Huang *et al.* were obtained from Gene Expression Omnibus (GEO), 2 replicates of each genotype at the 3-week timepoint. (WT: GSM2144682, GSM2144683; Rai1 cKO: GSM2144684, GSM2144685). Bulk RNA-seq data from Kasper *et al.* were obtained from ArrayExpress (62) (Accession: E-MTAB-10490, 4 replicates of each genotype). Mouse ESC and DIV7 neuron bulk RNA-seq data used in Fig. 7A was deposited to GEO, accession .

To identify differentiation and maturation genes, we performed PCA on WT samples and found that PC1 explained 78% of the variance. We then performed iterative PCA on genes with high loading on the initial PC1 (top 300, 500, and 1000 genes) and found that their PCA profiles were similar, suggesting the top 300 genes represent gene expression changes during differentiation of iNeurons. Hierarchical clustering of these core genes revealed two main clusters: cluster 1 shows continuous upregulation, while cluster 2 shows continuous downregulation (Fig. S7).

We then plotted the median z-scores of each cluster’s genes across genotypes and time points. At day 3, cluster 1 genes showed accelerated upregulation in both heterozygous and homozygous mutants. At day 7, cluster 1 genes showed continuous upregulation in WT, yet their expression began to taper down in the mutants—again in a *RAI1* dose-dependent manner. Meanwhile, cluster 2 genes showed a similar pattern of downregulation during differentiation across genotypes, with only a slight acceleration in the homozygous mutant. The number of genes chosen based on PCA loading did not change these results. These proof-of-principle analyses validated our strategy. Analysis was done in R. The number of centers for k-means clustering was chosen by the elbow method by plotting the within-cluster sum of squares.

### Immunoblotting

Immunoblotting was performed as previously described (21). 3×10⁵ cells were collected for each sample at day 10 of iNeuron differentiation. Cells were lysed, and proteins were extracted using RIPA buffer (Sigma R0278) following the manufacturer’s protocol. The extracted proteins were used for immunoblotting following the standard protocol. We used in-house rabbit polyclonal anti-RAI1 antibody (21) and anti-GAPDH (Cell Signaling D16H11).

### Electrophysiology

Miniature excitatory post-synaptic currents (mEPSC) were recorded as previously described with minor modification (63). Human neural progenitor cells (NPCs) were plated on coated Mattek dishes and maintained as described above. Recordings were performed at days 50-70 post-plating. Age-matched cultures from each genotype were recorded in a pairwise manner. Whole-cell voltage-clamp recordings were obtained from pyramidal-like neurons identified based on their morphology, characterized by a prominent apical dendrite and basal dendritic arborization. Patch pipettes with resistance of 3-6 MΩ were used. The internal solution contained (in mM): 100 Cs-gluconate, 0.2 EGTA, 5 MgCl2, 2 Mg-ATP, 0.3 Na-GTP and 40 HEPES (pH 7.20). The external solution contained (in mM): 119 NaCl, 5 KCl, 2 CaCl2, 2 MgCl2, 20 glucose, 10 HEPES. Tetrodotoxin (1µM) and bicuculline (40µM) were added to isolate mEPSCs. Cells were visualized using an Olympus IX-70 inverted microscope. Signals were amplified using a Multiclamp 700B (Axon Instruments) and digitized with a Digidata 1440 (Axon Instruments).

Voltage-clamp was performed at a Holding potential of -60 mV. Data were analyzed offline using Minianalysis (Synaptosoft). Typically, cells with an access resistance >25 M or with a >20% increase during recording were excluded from further analysis. Neurons exhibiting no detectable events during at least 1min of recording were also discarded. For statistical analysis, one-way ANOVA followed by Tukey’s post hoc tests were performed.

## Data availability

All plots are made with R, and scripts for all analyses are available on GitHub: <u>2025_rai1_hNeuron_public</u>. Sequencing data are available at NCBI Gene Expression Omnibus (GEO): GSE337485 (dual SMADi bulk RNA-seq), GSE337359 (scRNA-seq), GSE337750 (Mouse data), and GSE336167 (iNeuron RNA-seq),

## Acknowledgement

*RAI1*-deficient hESCs were generated by the University of Michigan Medical school Human Stem Cells and Gene Editing core Facility (RRID:SCR_026755). We thank the ENCODE Consortium for generating and organizing the data and making them accessible. Sequencing library preparation and next-generation sequencing were performed at the University of Michigan Advanced Genomics Core. Research reported in this publication was supported by the University of Michigan Advanced Genomics Core, the UM Single Cell Spatial Analysis Program, and the National Cancer Institute of the National Institutes of Health under Award Number P30CA046592 by the use of the following Cancer Center Shared Resource: Single Cell and Spatial Analysis Shared Resource. We thank Dr. Michael Uhler at the Michigan Neuroscience Institute for generating the iNeuron lines.

## Author contributions

BZ, MAS, and SI conceived the project. BZ validated hESC and iNeuron lines, performed dual SMADi differentiation. SM, PR, LTD, and SI analyzed sequencing data. TT performed electrophysiology experiments and analysis. GL were responsible for iNeurons differentiation. LTD, MAS, and SI acquired fundings. All authors wrote or edited the manuscript.

## Funding and additional information

The following funding supported the work: NIH/NINDS R21NS125449, R03NS137487, NS141983, MH133632 to SI Jim and Sandy Danto Family Research Fund. SMS Research Foundation Grant (to MAS and SI).

---

**Figure S2:**
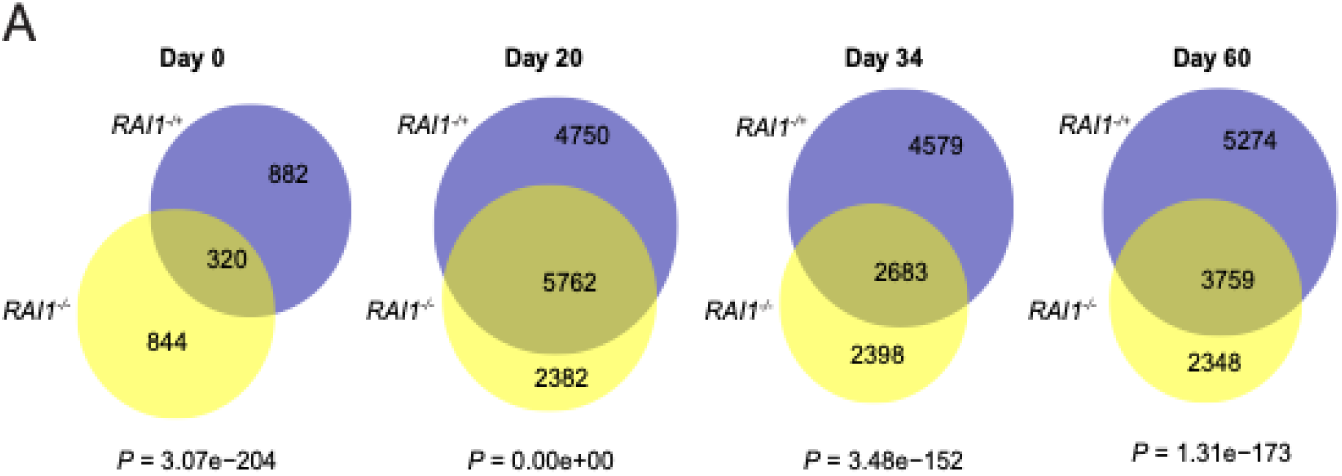
Differentially expressed genes overlap between *RAI1*+/- and *RAI1*-/-cells. **(A)** The number of DEGs (Padj < 0.05) in *RAI1*+/- and *RAI1*-/-compared to WT controls was denoted, and their identities were compared. P-values of the chi-square test are denoted at the bottom of the Venn diagrams.

**Figure S5:**
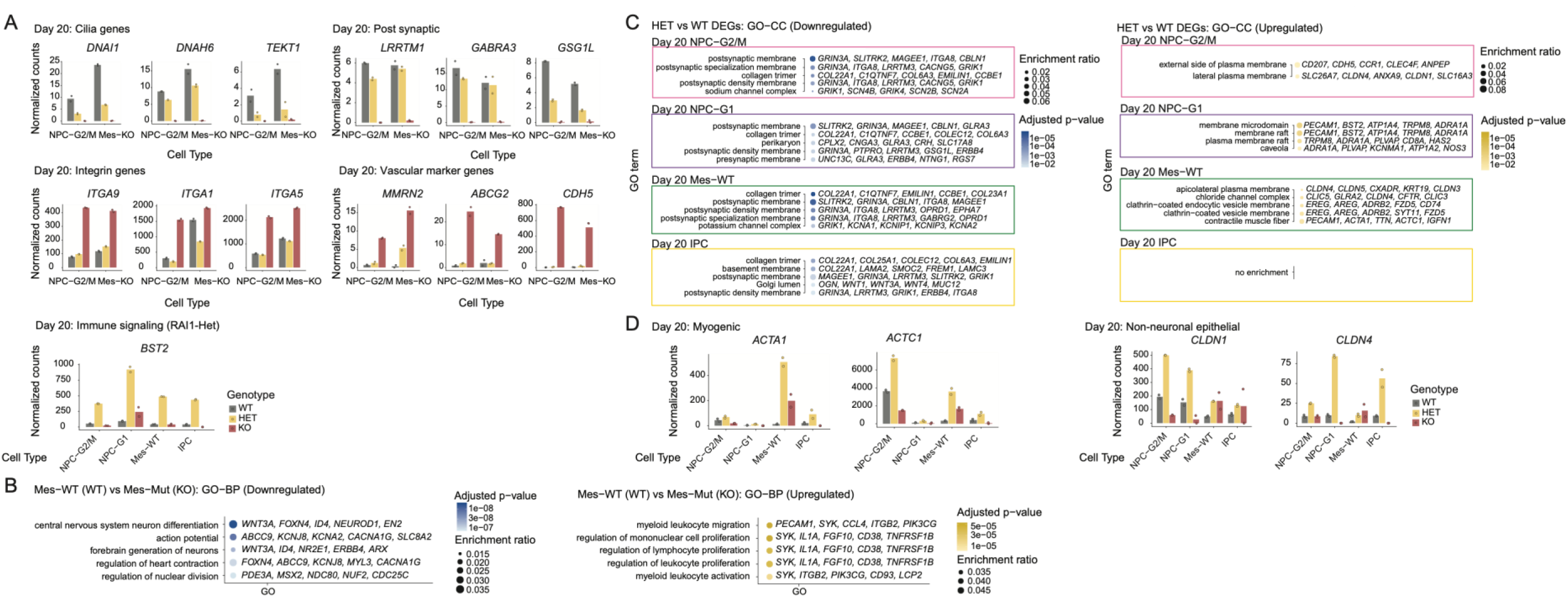
Additional analyses of genes dysregulated in *RAI1*-deficient cell types on day 20. **(A)** Expression of additional top contributors for GO term enrichment. **(B)** GO Cellular Component enrichment for *RAI1*-heterozygous DEGs. The top five GO terms are presented with the genes that contributed most to the enrichment. **(C)** GO Biological Processes enrichment analysis comparing Mes-WT (WT) and Mes-KO (KO). **(D)** Expression of the additional myogenic and non-neuronal epithelial genes across genotypes and cell types. Mean values are shown in histograms, with individual data points from biological duplicates.

**Figure S6:**
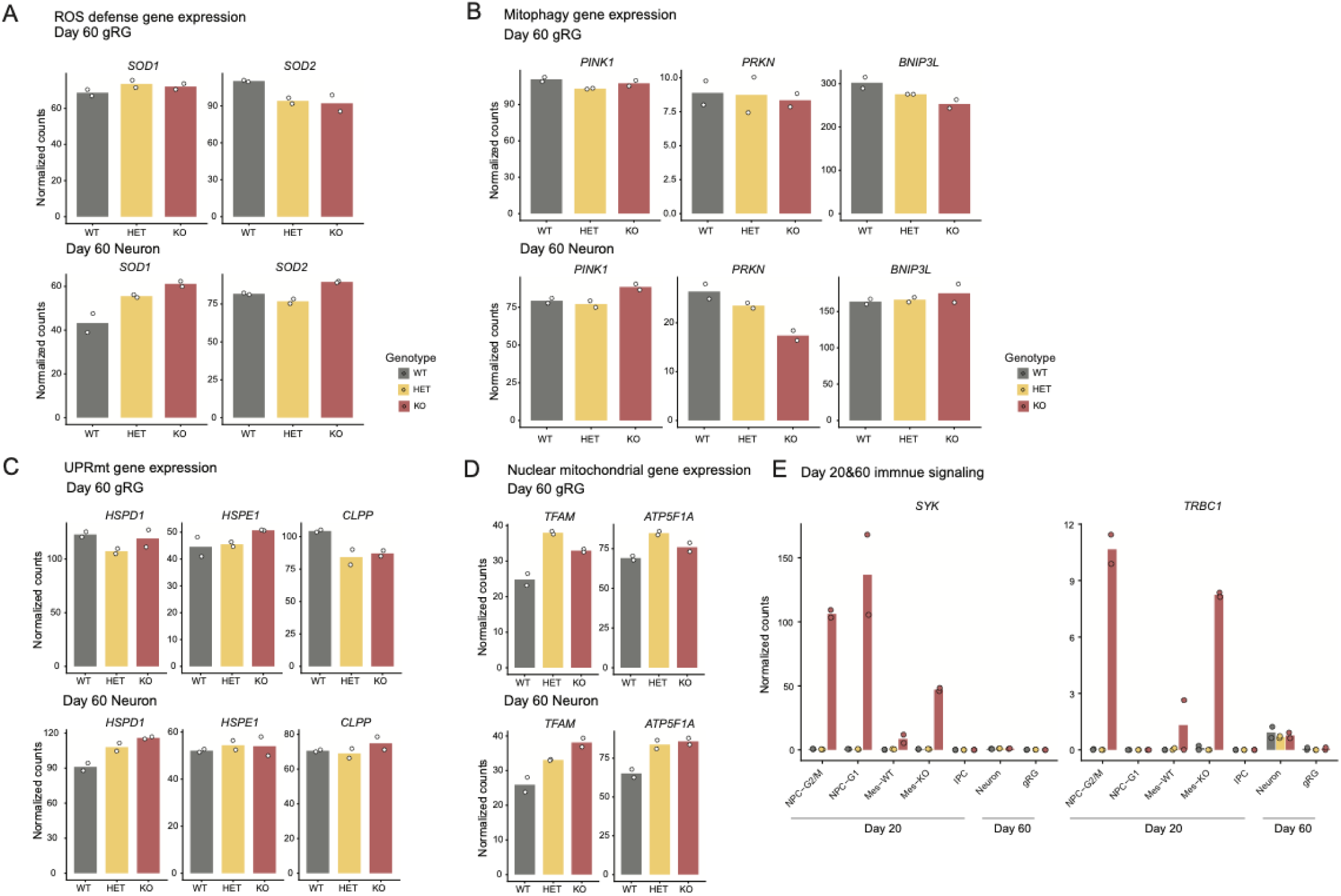
Additional gene expression plots of *RAI1*-deficient cell types on day 60. **(A)** Expression of ROS scavengers. **(B)** Expression of mitophagy-related genes. **(C)** Expression of genes related to unfolded protein responses at mitochondria (UPRmt). **(D)** Expression of nuclear genome-encoded respiratory change regulator genes. Mean values are shown in histograms, with individual data points from biological duplicates. **(E)** Additional immune signaling genes were derepressed only in day 20 cells.

**Figure S7:**
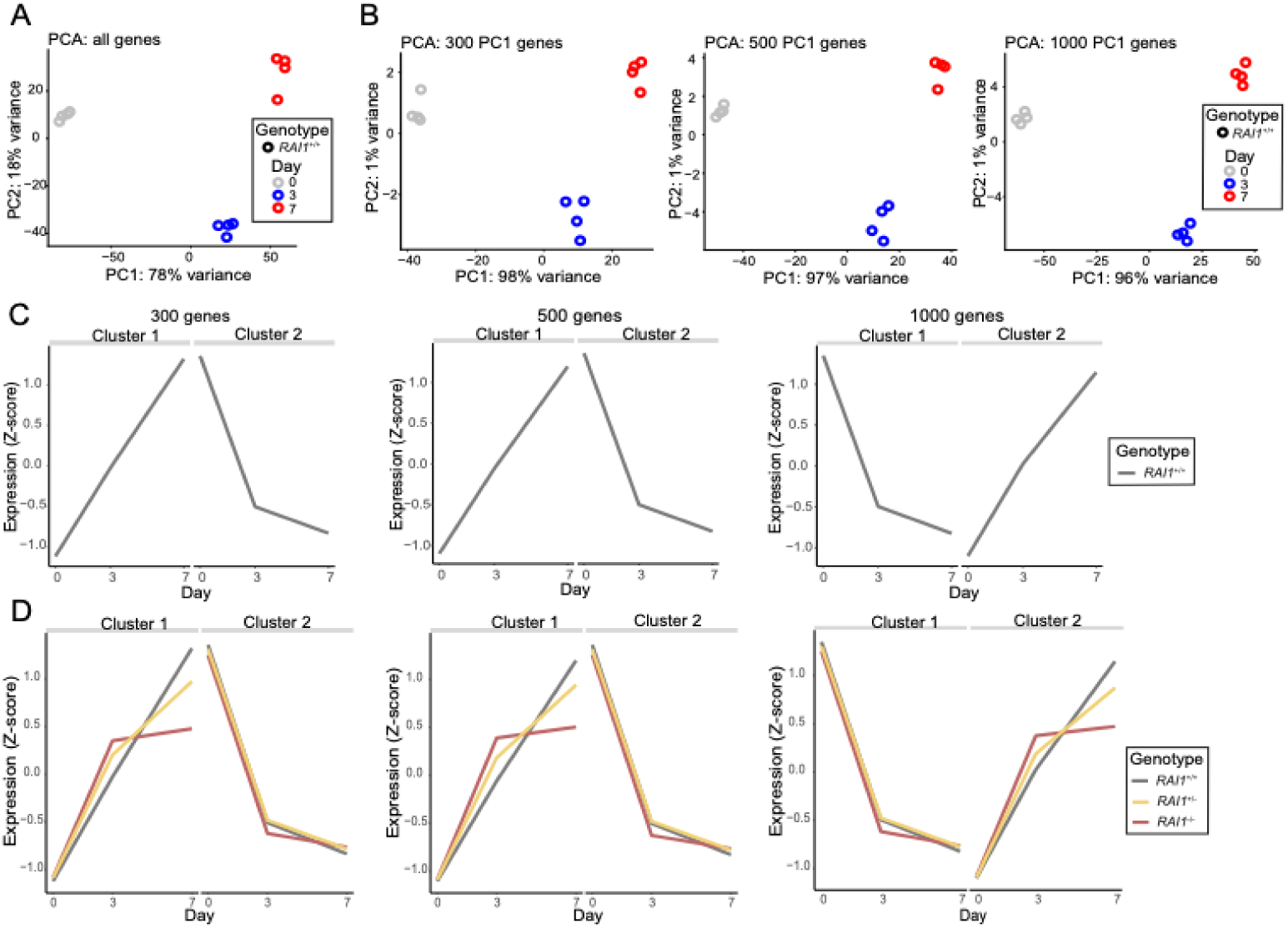
Principal component identification of core *Rai1*-dependent accelerated genes in iNeurons. **(A)** PCA of all genes in WT iNeurons. **(B)** PCA using the top 300, 500, and 1000 genes with the largest absolute gene loadings on PC1 from the plot in part A. **(C)** K-means clustering of gene expression in the top 300, 500, and 1000 genes with the greatest absolute value of gene loading on PC1 identified in part A. **(D)** Gene expression of WT, Het, and KO iNeurons within each cluster, identified from the corresponding number of top PC1 gene loadings shown in part B.

**Fig S8:**
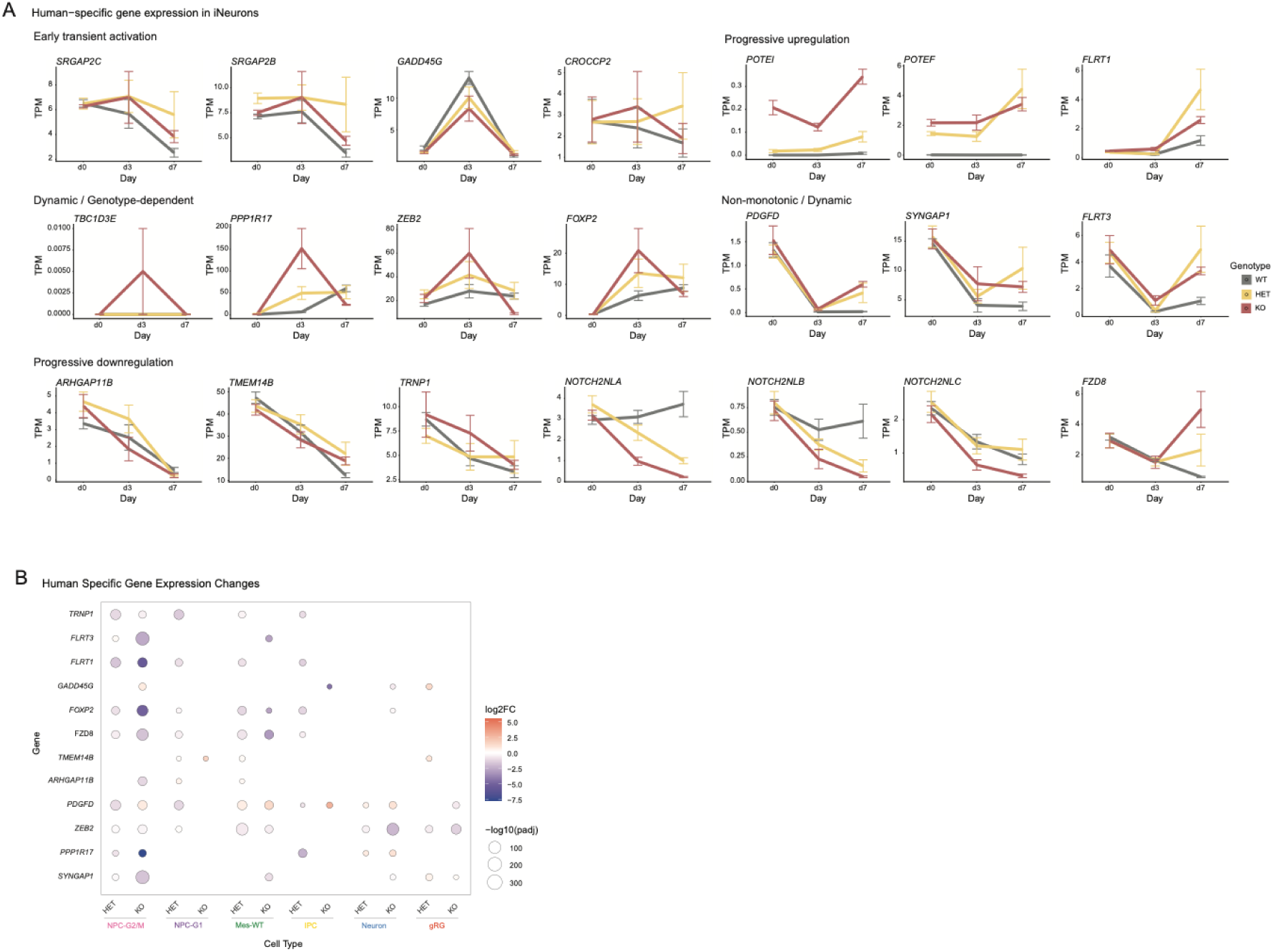
Expression of primate/human-specific genes implicated in the slower tempo of brain development. **(A)** The genes are categorized based on how their expression changes in WT human iNeurons. **(B)** Expression of human primate-specific genes in single-cell RNAseq data. These genes are covered by the 10x Chromium Fixed RNA Profiling (Flex) genomic probes.

## Notes

### Competing Interest Statement

The authors have declared no competing interest.

### Summary of Updates

The texts are edited for accuracy. The figures are enlarged for easier viewing.

## References

1. Wallace JL, Pollen AA. Human neuronal maturation comes of age: cellular mechanisms and species differences. Nat Rev Neurosci. 2024 Jan;25(1):7–29. doi:10.1038/s41583-023-00760-3

2. Vanderhaeghen P, Polleux F. Developmental mechanisms underlying the evolution of human cortical circuits. Nat Rev Neurosci. 2023 Apr;24(4):213–32. doi:10.1038/s41583-023-00675-z

3. Sherwood CC, Gómez-Robles A. Brain Plasticity and Human Evolution. Annu Rev Anthropol. 2017 Oct 23;46(1):399–419. doi:10.1146/annurev-anthro-102215-100009

4. Bae BI, Jayaraman D, Walsh CA. Genetic Changes Shaping the Human Brain. Dev Cell. 2015 Feb;32(4):423–34. doi:10.1016/j.devcel.2015.01.035

5. Benito-Kwiecinski S, Giandomenico SL, Sutcliffe M, Riis ES, Freire-Pritchett P, Kelava I, et al. An early cell shape transition drives evolutionary expansion of the human forebrain. Cell. 2021 Apr;184(8):2084–2102.e19. doi:10.1016/j.cell.2021.02.050

6. Suzuki IK, Gacquer D, Van Heurck R, Kumar D, Wojno M, Bilheu A, et al. Human-Specific NOTCH2NL Genes Expand Cortical Neurogenesis through Delta/Notch Regulation. Cell. 2018 May;173(6):1370–1384.e16. doi:10.1016/j.cell.2018.03.067

7. Iwata R, Vanderhaeghen P. Metabolic mechanisms of species-specific developmental tempo. Dev Cell. 2024 Jul;59(13):1628–39. doi:10.1016/j.devcel.2024.05.027

8. Ciceri G, Baggiolini A, Cho HS, Kshirsagar M, Benito-Kwiecinski S, Walsh RM, et al. An epigenetic barrier sets the timing of human neuronal maturation. Nature. 2024 Feb 22;626(8000):881–90. doi:10.1038/s41586-023-06984-8

9. Iossifov I, O’Roak BJ, Sanders SJ, Ronemus M, Krumm N, Levy D, et al. The contribution of de novo coding mutations to autism spectrum disorder. Nature. 2014 Nov;515(7526):216–21. doi:10.1038/nature13908

10. De Rubeis S, He X, Goldberg AP, Poultney CS, Samocha K, Cicek AE, et al. Synaptic, transcriptional, and chromatin genes disrupted in autism. Nature. 2014 Nov 13;515(7526):209–15. doi:10.1038/nature13772 PubMed PMID: 25363760; PubMed Central PMCID: PMC4402723.

11. Network and Pathway Analysis Subgroup of Psychiatric Genomics Consortium. Psychiatric genome-wide association study analyses implicate neuronal, immune and histone pathways. Nat Neurosci. 2015 Feb;18(2):199–209. doi:10.1038/nn.3922 PubMed PMID: 25599223; PubMed Central PMCID: PMC4378867.

12. Slager RE, Newton TL, Vlangos CN, Finucane B, Elsea SH. Mutations in RAI1 associated with Smith–Magenis syndrome. Nat Genet. 2003 Apr;33(4):466–8. doi:10.1038/ng1126

13. Smith AC, Berens J, Boyd KE, Brennan C, Gropman A, Haas-Givler B, et al. Smith-Magenis Syndrome. In: Adam MP, Bick S, Mirzaa GM, Pagon RA, Wallace SE, Amemiya A, editors. GeneReviews® [Internet]. Seattle (WA): University of Washington, Seattle; 1993 [cited 2026 Jun 4]. Available from: http://www.ncbi.nlm.nih.gov/books/NBK1310/ PubMed PMID: 20301487.

14. Girirajan S, Vlangos CN, Szomju BB, Edelman E, Trevors CD, Dupuis L, et al. Genotype–phenotype correlation in Smith-Magenis syndrome: Evidence that multiple genes in 17p11.2 contribute to the clinical spectrum. Genet Med. 2006 Jul;8(7):417–27. doi:10.1097/01.gim.0000228215.32110.89

15. Bi W, Ohyama T, Nakamura H, Yan J, Visvanathan J, Justice MJ, et al. Inactivation of Rai1 in mice recapitulates phenotypes observed in chromosome engineered mouse models for Smith–Magenis syndrome. Hum Mol Genet. 2005 Apr 15;14(8):983–95. doi:10.1093/hmg/ddi085

16. Bi W, Yan J, Shi X, Yuva-Paylor LA, Antalffy BA, Goldman A, et al. Rai1 deficiency in mice causes learning impairment and motor dysfunction, whereas Rai1 heterozygous mice display minimal behavioral phenotypes. Hum Mol Genet. 2007 Aug 1;16(15):1802–13. doi:10.1093/hmg/ddm128

17. Burns B, Schmidt K, Williams SR, Kim S, Girirajan S, Elsea SH. *Rai1* haploinsufficiency causes reduced *Bdnf* expression resulting in hyperphagia, obesity and altered fat distribution in mice and humans with no evidence of metabolic syndrome. Hum Mol Genet. 2010 Oct 15;19(20):4026–42. doi:10.1093/hmg/ddq317

18. Huang WH, Guenthner CJ, Xu J, Nguyen T, Schwarz LA, Wilkinson AW, et al. Molecular and Neural Functions of Rai1, the Causal Gene for Smith-Magenis Syndrome. Neuron. 2016 Oct;92(2):392–406. doi:10.1016/j.neuron.2016.09.019

19. Lee YJ, Chang YT, Cho Y, Kowalczyk M, Dragoiescu A, Pacis A, et al. Molecular and developmental deficits in Smith-Magenis syndrome human stem cell-derived cortical neural models. Am J Hum Genet. 2025 Oct;112(10):2338–62. doi:10.1016/j.ajhg.2025.07.020

20. Supek F, Lehner B, Lindeboom RGH. To NMD or Not To NMD: Nonsense-Mediated mRNA Decay in Cancer and Other Genetic Diseases. Trends Genet. 2021 Jul;37(7):657–68. doi:10.1016/j.tig.2020.11.002

21. Garay PM, Chen A, Tsukahara T, Rodríguez Díaz JC, Kohen R, Althaus JC, et al. RAI1 Regulates Activity-Dependent Nascent Transcription and Synaptic Scaling. Cell Rep. 2020 Aug;32(6):108002. doi:10.1016/j.celrep.2020.108002

22. Yang T, Pang D, Li C, Shang H. Retinoic Acid-Induced 1 Gene and Neuropsychiatric Diseases: A Systematic Review. Expert Rev Mol Med. 2025 May 29;27:e17. doi:10.1017/erm.2025.12 PubMed PMID: 40437981; PubMed Central PMCID: PMC12133160.

23. Chambers SM, Fasano CA, Papapetrou EP, Tomishima M, Sadelain M, Studer L. Highly efficient neural conversion of human ES and iPS cells by dual inhibition of SMAD signaling. Nat Biotechnol. 2009 Mar;27(3):275–80. doi:10.1038/nbt.1529

24. Satija R, Farrell JA, Gennert D, Schier AF, Regev A. Spatial reconstruction of single-cell gene expression data. Nat Biotechnol. 2015 May;33(5):495–502. doi:10.1038/nbt.3192

25. Menendez L, Yatskievych TA, Antin PB, Dalton S. Wnt signaling and a Smad pathway blockade direct the differentiation of human pluripotent stem cells to multipotent neural crest cells. Proc Natl Acad Sci. 2011 Nov 29;108(48):19240–5. doi:10.1073/pnas.1113746108

26. Trapnell C, Cacchiarelli D, Grimsby J, Pokharel P, Li S, Morse M, et al. The dynamics and regulators of cell fate decisions are revealed by pseudotemporal ordering of single cells. Nat Biotechnol. 2014 Apr;32(4):381–6. doi:10.1038/nbt.2859

27. Squair JW, Gautier M, Kathe C, Anderson MA, James ND, Hutson TH, et al. Confronting false discoveries in single-cell differential expression. Nat Commun. 2021 Sep 28;12(1):5692. doi:10.1038/s41467-021-25960-2

28. Buckley CE, St Johnston D. Apical–basal polarity and the control of epithelial form and function. Nat Rev Mol Cell Biol. 2022 Aug;23(8):559–77. doi:10.1038/s41580-022-00465-y

29. Gudjohnsen SAH, Atacho DAM, Gesbert F, Raposo G, Hurbain I, Larue L, et al. Meningeal Melanocytes in the Mouse: Distribution and Dependence on Mitf. Front Neuroanat. 2015 Nov 25;9. doi:10.3389/fnana.2015.00149

30. Ando K, Tong L, Peng D, Vázquez-Liébanas E, Chiyoda H, He L, et al. KCNJ8/ABCC9-containing K-ATP channel modulates brain vascular smooth muscle development and neurovascular coupling. Dev Cell. 2022 Jun;57(11):1383–1399.e7. doi:10.1016/j.devcel.2022.04.019

31. Azevedo PO, Sena IFG, Andreotti JP, Carvalho-Tavares J, Alves-Filho JC, Cunha TM, et al. Pericytes modulate myelination in the central nervous system. J Cell Physiol. 2018 Aug;233(8):5523–9. doi:10.1002/jcp.26348

32. Marques F, Sousa JC, Coppola G, Falcao AM, Rodrigues AJ, Geschwind DH, et al. Kinetic Profile of the Transcriptome Changes Induced in the Choroid Plexus by Peripheral Inflammation. J Cereb Blood Flow Metab. 2009 May;29(5):921–32. doi:10.1038/jcbfm.2009.15

33. Zheng X, Boyer L, Jin M, Mertens J, Kim Y, Ma L, et al. Metabolic reprogramming during neuronal differentiation from aerobic glycolysis to neuronal oxidative phosphorylation. eLife. 2016 Jun 10;5:e13374. doi:10.7554/eLife.13374

34. A O, U M, Lf B, A GC. Energy metabolism in childhood neurodevelopmental disorders. eBioMedicine. 2021 Jul;69:103474. doi:10.1016/j.ebiom.2021.103474

35. 35. Käsper EL, Hwang IY, Grötsch H, Fung HKH, Sérandour AA, Arecco N, et al. The neurodevelopmental disorder-linked PHF14 complex that forms biomolecular condensates detects DNA damage and promotes repair [Internet]. bioRxiv; 2021 [cited 2026 Jun 18]. p. 2021.10.12.462922. Available from: https://www.biorxiv.org/content/10.1101/2021.10.12.462922v1 doi:10.1101/2021.10.12.462922

36. Zhang Y, Pak C, Han Y, Ahlenius H, Zhang Z, Chanda S, et al. Rapid Single-Step Induction of Functional Neurons from Human Pluripotent Stem Cells. Neuron. 2013 Jun;78(5):785–98. doi:10.1016/j.neuron.2013.05.029

37. Gupta S, M-Redmond T, Meng F, Tidball A, Akil H, Watson S, et al. Fibroblast growth factor 2 regulates activity and gene expression of human post-mitotic excitatory neurons. J Neurochem. 2018 May;145(3):188–203. doi:10.1111/jnc.14255

38. Libé-Philippot B, Vanderhaeghen P. Cellular and Molecular Mechanisms Linking Human Cortical Development and Evolution. Annu Rev Genet. 2021 Nov 23;55(1):555–81. doi:10.1146/annurev-genet-071719-020705

39. Fiddes IT, Lodewijk GA, Mooring M, Bosworth CM, Ewing AD, Mantalas GL, et al. Human-Specific NOTCH2NL Genes Affect Notch Signaling and Cortical Neurogenesis. Cell. 2018 May;173(6):1356–1369.e22. doi:10.1016/j.cell.2018.03.051

40. Nagao M, Lanjakornsiripan D, Itoh Y, Kishi Y, Ogata T, Gotoh Y. High Mobility Group Nucleosome-Binding Family Proteins Promote Astrocyte Differentiation of Neural Precursor Cells. Stem Cells. 2014 Nov 1;32(11):2983–97. doi:10.1002/stem.1787

41. Yang YJ, Baltus AE, Mathew RS, Murphy EA, Evrony GD, Gonzalez DM, et al. Microcephaly Gene Links Trithorax and REST/NRSF to Control Neural Stem Cell Proliferation and Differentiation. Cell. 2012 Nov;151(5):1097–112. doi:10.1016/j.cell.2012.10.043

42. Wang H, Ge G, Uchida Y, Luu B, Ahn S. *Gli3* Is Required for Maintenance and Fate Specification of Cortical Progenitors. J Neurosci. 2011 Apr 27;31(17):6440–8. doi:10.1523/JNEUROSCI.4892-10.2011

43. Shimada IS, Acar M, Burgess RJ, Zhao Z, Morrison SJ. Prdm16 is required for the maintenance of neural stem cells in the postnatal forebrain and their differentiation into ependymal cells. Genes Dev. 2017 Jun 1;31(11):1134–46. doi:10.1101/gad.291773.116

44. DeSisto J, O’Rourke R, Jones HE, Pawlikowski B, Malek AD, Bonney S, et al. Single-Cell Transcriptomic Analyses of the Developing Meninges Reveal Meningeal Fibroblast Diversity and Function. Dev Cell. 2020 Jul;54(1):43–59.e4. doi:10.1016/j.devcel.2020.06.009

45. Lacaria M, Gu W, Lupski JR. Circadian abnormalities in mouse models of smith–magenis syndrome: Evidence for involvement of *RAI1*. Am J Med Genet A. 2013 Jul;161(7):1561–8. doi:10.1002/ajmg.a.35941

46. Huang WH, Wang DC, Allen WE, Klope M, Hu H, Shamloo M, et al. Early adolescent Rai1 reactivation reverses transcriptional and social interaction deficits in a mouse model of Smith–Magenis syndrome. Proc Natl Acad Sci. 2018 Oct 16;115(42):10744–9. doi:10.1073/pnas.1806796115

47. Yan J, Bi W, Lupski JR. Penetrance of Craniofacial Anomalies in Mouse Models of Smith-Magenis Syndrome Is Modified by Genomic Sequence Surrounding Rai1: Not All Null Alleles Are Alike. Am J Hum Genet. 2007 Mar;80(3):518–25. doi:10.1086/512043

48. Tahir R, Kennedy A, Elsea SH, Dickinson AJ. Retinoic acid induced-1 (Rai1) regulates craniofacial and brain development in Xenopus. Mech Dev. 2014 Aug;133:91–104. doi:10.1016/j.mod.2014.05.004

49. Pendleton AL, Shen F, Taravella AM, Emery S, Veeramah KR, Boyko AR, et al. Comparison of village dog and wolf genomes highlights the role of the neural crest in dog domestication. BMC Biol. 2018 Dec;16(1):64. doi:10.1186/s12915-018-0535-2

50. 50. Zhang Lab. CRISPR Design Tool [Internet]. Genome Engineering; [cited 2026 Jun 5]. Available from: https://www.genome-engineering.org/crispr/

51. Ran FA, Hsu PD, Wright J, Agarwala V, Scott DA, Zhang F. Genome engineering using the CRISPR-Cas9 system. Nat Protoc. 2013 Nov;8(11):2281–308. doi:10.1038/nprot.2013.143

52. Agarwal S, Bonefas KM, Garay PM, Brookes E, Murata-Nakamura Y, Porter RS, et al. KDM1A maintains genome-wide homeostasis of transcriptional enhancers. Genome Res. 2021 Feb;31(2):186–97. doi:10.1101/gr.234559.118

53. Bonefas KM, Vallianatos CN, Raines B, Tronson NC, Iwase S. Sexually Dimorphic Alterations in the Transcriptome and Behavior with Loss of Histone Demethylase KDM5C. Cells. 2023 Feb 16;12(4):637. doi:10.3390/cells12040637

54. Köster J, Rahmann S. Snakemake—a scalable bioinformatics workflow engine. Bioinformatics. 2012 Oct 1;28(19):2520–2. doi:10.1093/bioinformatics/bts480

55. Martin M. Cutadapt removes adapter sequences from high-throughput sequencing reads. EMBnet.journal. 2011 May 2;17(1):10. doi:10.14806/ej.17.1.200

56. Dobin A, Davis CA, Schlesinger F, Drenkow J, Zaleski C, Jha S, et al. STAR: ultrafast universal RNA-seq aligner. Bioinformatics. 2013 Jan 1;29(1):15–21. doi:10.1093/bioinformatics/bts635

57. Li B, Dewey CN. RSEM: accurate transcript quantification from RNA-Seq data with or without a reference genome. BMC Bioinformatics. 2011 Dec;12(1):323. doi:10.1186/1471-2105-12-323

58. Ewels P, Magnusson M, Lundin S, Käller M. MultiQC: summarize analysis results for multiple tools and samples in a single report. Bioinformatics. 2016 Oct 1;32(19):3047–8. doi:10.1093/bioinformatics/btw354

59. The ENCODE Project Consortium. An integrated encyclopedia of DNA elements in the human genome. Nature. 2012 Sep;489(7414):57–74. doi:10.1038/nature11247

60. Kagda MS, Lam B, Litton C, Small C, Sloan CA, Spragins E, et al. Data navigation on the ENCODE portal. Nat Commun. 2025 Oct 30;16(1):9592. doi:10.1038/s41467-025-64343-9

61. 61. ENCODE [Internet]. [cited 2026 Jun 4]. Available from: https://www.encodeproject.org/

62. Parkinson H, Kapushesky M, Shojatalab M, Abeygunawardena N, Coulson R, Farne A, et al. ArrayExpress--a public database of microarray experiments and gene expression profiles. Nucleic Acids Res. 2007 Jan;35(Database issue):D747-750. doi:10.1093/nar/gkl995 PubMed PMID: 17132828; PubMed Central PMCID: PMC1716725.

63. Tsukahara T, Kethireddy S, Bonefas KM, Chen A, Sutton BLM, Bandow K, et al. Division of labor among H3K4 methyltransferases defines distinct facets of homeostatic plasticity. Cell Rep. 2025 Jun;44(6):115746. doi:10.1016/j.celrep.2025.115746

